# The retrotransposon*-*derived capsid genes *PNMA1* and *PNMA4* maintain reproductive capacity

**DOI:** 10.1101/2024.05.11.592987

**Authors:** Thomas W.P. Wood, William S. Henriques, Harrison B. Cullen, Mayra Romero, Cecilia S. Blengini, Shreya Sarathy, Julia Sorkin, Hilina Bekele, Chen Jin, Seungsoo Kim, Alexei Chemiakine, Rishad C. Khondker, José V.V. Isola, Michael B. Stout, Vincenzo A. Gennarino, Binyam Mogessie, Devanshi Jain, Karen Schindler, Yousin Suh, Blake Wiedenheft, Luke E. Berchowitz

**Author notes:** Correspondence; Luke E. Berchowitz, Ph.D., Columbia University Irving Medical Center, 701 W 168th St. Hammer Health Sciences Building Room 1520, New York NY, 10032.

## Abstract

The human genome contains 24 *gag*-like capsid genes derived from deactivated retrotransposons conserved among eutherians. Although some of their encoded proteins retain the ability to form capsids and even transfer cargo, their fitness benefit has remained elusive. Here we show that the *gag*-like genes *PNMA1* and *PNMA4* support reproductive capacity. Six-week-old mice lacking either *Pnma1* or *Pnma4* are indistinguishable from wild-type littermates, but by six months the mutant mice become prematurely subfertile, with precipitous drops in sex hormone levels, gonadal atrophy, and abdominal obesity; overall they produce markedly fewer offspring than controls. Analysis of donated human ovaries shows that expression of both genes declines normally with aging, while several *PNMA1* and *PNMA4* variants identified in genome-wide association studies are causally associated with low testosterone, altered puberty onset, or obesity. These findings expand our understanding of factors that maintain human reproductive health and lend insight into the domestication of retrotransposon-derived genes.

## INTRODUCTION

Almost half of the human genome consists of retrotransposons, ‘parasitic’ sequences that insert themselves into the host genome via an RNA intermediate^1^. While most of these sequences are silenced or mutationally deactivated, they can present opportunities for evolutionary innovation: mutation of a deteriorating retrotransposon can, in rare circumstances, result in a gene that provides a selective advantage to the host in a process termed domestication^2–5^. In order for a domesticated gene to be beneficial, however, it needs to acquire a useful expression pattern. To maintain vertical inheritance and avoid extinction, a retrotransposon must express and proliferate in germ cells or their precursors^6–8^. This is no trivial feat, as host cells have numerous strategies to silence retrotransposon sequences including DNA methylation, chromatin modification, and transcriptional interference^7,9^. It is possible to evade these maneuvers, however, as the yeast *Metaviridae* (formerly *Ty3*) long-terminal repeat (LTR) retrotransposons do, by integrating downstream of an essential meiotic transcription factor binding site during germ cell development^10^. Could this evolutionary contest between host and retrotransposon shape the activities of the domesticated genes?

The *PNMA* family of *gag*-like genes, which were domesticated from an ancient vertebrate retrotransposon of the *Metaviridae* clade at least 100 million years ago^11,12^, may provide an affirmative answer to that question. There are six full-length *PNMA* genes in humans and five in mice^13^. PNMA1-3 were originally identified as targets of the autoantibodies that cause paraneoplastic neurological syndrome (PNS) in lung and gynecologic cancers. *PNMA4-6* were identified based on homology with *PNMA1-3* but are not implicated in PNS^14–16^. We reasoned that the conservation of *PNMA* genes is unlikely to be based on their autoantigenic properties after ectopic expression in neoplasms. While *Pnma5* is implicated in mouse oocyte development via an unknown mechanism^17^, little is known about the non-pathological roles of the other genes other than that some seem to have retained the ability to form capsids when expressed in human cells or bacteria^18,19^. The autoantigenicity of PNMA2, in fact, requires capsid formation^20^. *PNMA1* and *PNMA4* (also called *MOAP-1* or *MAP-1*) harbor capsid-like, linker, and N-terminal RNA-binding domains (RBDs)^12,13^, though whether they form capsids *in situ* is unknown. They are, however, positively regulated by the master germ cell transcription factors MYBL1 and STRA8, and their transcripts are bound by the translational regulator DAZL during gametogenesis^10^. This developmental regulation of *PNMA1* and *PNMA4* expression in male and female gonadal tissue suggested to us that they might serve a reproductive function. Further support for this notion comes from the fact that these two genes co-reside within a syntenic block that has persisted throughout the evolution of placental mammals (∼20 mb apart on chromosome 14 in humans and ∼20 mb apart on chromosome 12 in mice). Given that marsupials also harbor a single *PNMA* (*mPNMA*) that is highly expressed in pachytene spermatocytes^11,21^, the germ cell expression of *PNMA* family genes was either present in the common therian ancestor or convergently evolved in marsupials and eutherians.

We therefore set out to determine whether either *PNMA1* or *PNMA4* affect reproductive function. Through the analysis of genetic mouse models, donated human ovaries, and genome-wide association studies (GWAS), we find that PNMA proteins are necessary for the maintenance of a normal fertility span. PNMA1 and PNMA4 self-assemble into capsids that exit human cells, and we observed large PNMA4 particles in male mouse gonadal tissue that are consistent with capsid formation.

## RESULTS

### Male and female mice lacking *Pnma1* or *Pnma4* show premature loss of fertility

*PNMA1* is universally conserved among eutherians and exhibits a purifying selection signature (**Figure 1A**, dN/dS = 0.06)^12^. The *PNMA4* locus is also universal to eutherians but has undergone more diversification: some lineages have acquired a premature stop within the coding sequence leading to pseudogenization and, in primates, an Alu element has integrated in the 3’ UTR. Despite its pseudogenization in some mammals, *PNMA4* also shows evidence of purifying selection (**Figure 1B**, dN/dS = 0.29)^12^. If *PNMA1* and *PNMA4* have been retained in mammals for reproductive functions, then we expect that animals lacking one or more of these genes should show some sort of diminution in reproductive capacity. We therefore used CRISPR-Cas9 with flanking cut sites to generate *Pnma1^-/-^* and *Pnma4^-/-^* mouse knockouts (KOs, **Figure S1A**), and we verified that both null alleles exhibited Mendelian segregation. To validate the knockouts and to assess whether the two genes compensate for one another, we analyzed *Pnma1* and *Pnma4* mRNA and protein in wild type and knockouts. Rather than seeing loss of one gene leading to compensatory upregulation of the other, we observed post-transcriptional anticompensation in both knockouts: *Pnma1^-/-^* testes showed an >80% reduction in PNMA4 protein levels (but unaltered mRNA), and *vice versa* for *Pnma4^-/-^* (**Figure S1B, C**).

**Figure 1:**
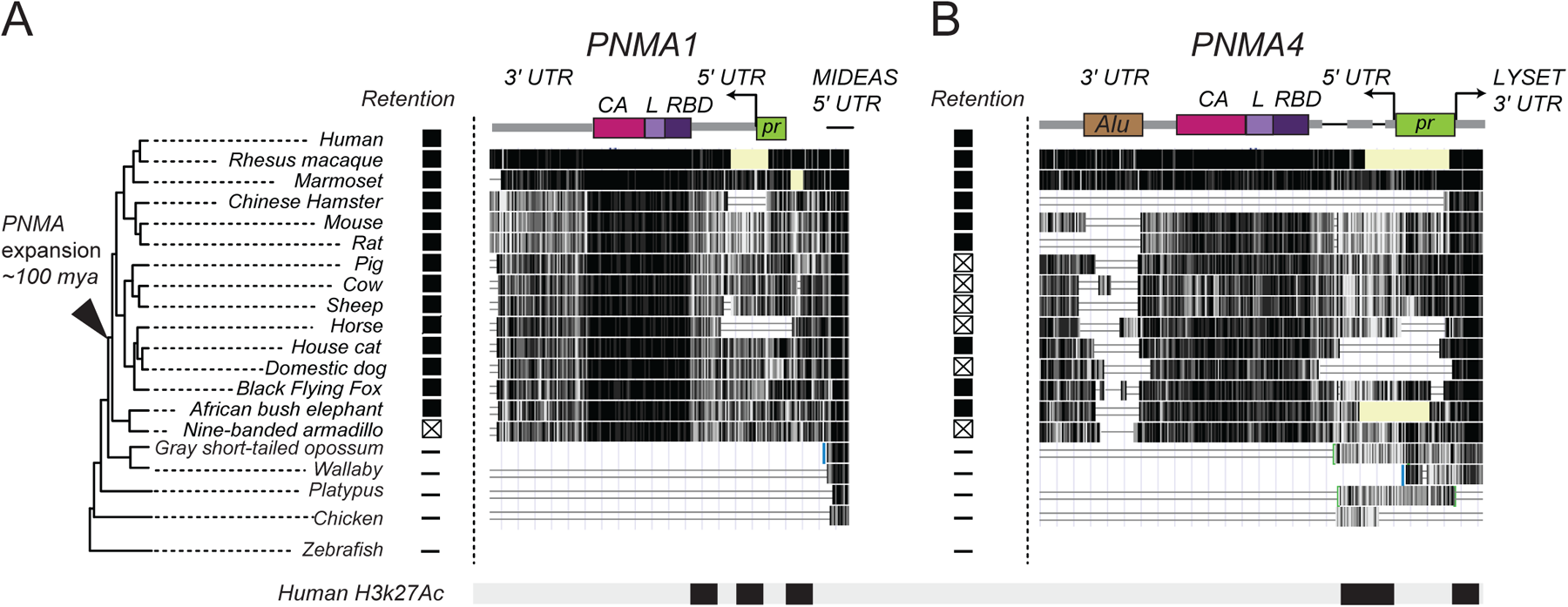
*PNMA1* and *PNMA4* are conserved among eutherians. **(A, B)** Architecture of the human *PNMA1* and *PNMA4*. Loci are colored according to domains: capsid domain (CA, magenta), linker (L, lilac), and RNA-binding domain (RBD, purple). The promoter (green) and transcription start sites are shown using arrows. A black triangle on the phylogenetic tree (left) indicates the point of the first expansion of the ancestral *PNMA* locus leading to *PNMA1-5*. *PNMA1* is universally retained across placental mammals as an intact gene, whereas *PNMA4* has experienced lineage-specific pseudogenization (boxed x’s). Conservation at each amino acid is shown with a vertical black line for fifteen eutherian mammals, three marsupials, and two vertebrate outgroups. The histone modification for active transcription (H3K27Ac) in humans is shown along the bottom.

The male knockout mice did not differ from wild-type littermates at two months of age, but very soon their fertility declined, as evidenced by litter size (**Figure 2A**): at three months of age, the mutants fathered litters ∼70% of the size of wild-type mice, and by six months of age, their litters were only 25-50% as large as those sired by wild-type mice (**Figure 2A, S1D**). There was a concomitant age-dependent testicular atrophy and decline in epididymal sperm count in single and double mutants compared to controls (**Figures 2B-D**). Serum testosterone levels in *Pnma1^-/-^* were statistically indistinguishable from controls at 1.5 and 3 months but fell precipitously by 6 months of age (**Figure 2E**). *Pnma4^-/-^* and double knockout mice showed less than 10% of the serum testosterone levels found in control animals at any time point. Even though peritubular myoid cells (which help transport sperm) and Sertoli cells (which support the development of immature sperm) appeared normal, from 3 months onward, some seminiferous tubules in single and double mutants were devoid of undifferentiated spermatogonia, spermatocytes, and spermatids (**Figures 2F, G, S2A**).

**Figure 2:**
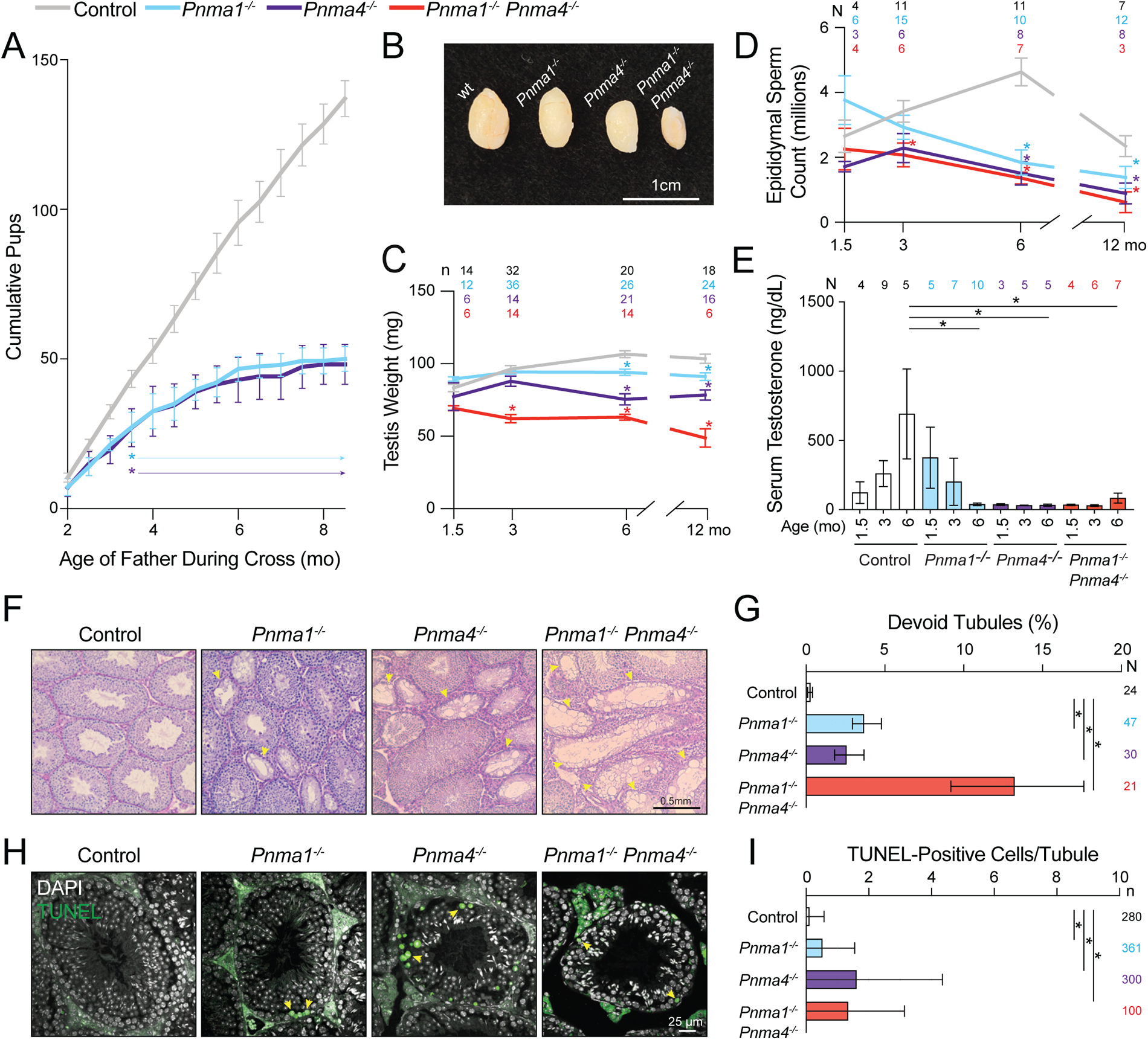
Male mice lacking *Pnma1* or *Pnma4* prematurely lose reproductive capacity. **(A)** Wild type male control mice (gray), *Pnma1^-/-^* (cyan), or *Pnma4^-/-^* (purple) were crossed biweekly to CF-1 female fertility tester mice (N = 5 pairs for each genotype-timepoint combination). Pup numbers for each cross were recorded. **(B, C)** Testes (n above graph) from control (pooled wild type and heterozygous, gray), *Pnma1^-/-^* (cyan), *Pnma4^-/-^* (purple), or *Pnma1^-/-^ Pnma4^-/-^* double mutant (red) mice were weighed. Representative images and quantifications are shown. **(D)** Sperm counts from dissected cauda epididymides. The number of individuals (N) for each genotype and age are shown above. **(E)** Serum testosterone (from N individual males) at the indicated ages. **(F)** PAS-stained tubule sections from 6-month-old mouse testes; devoid tubules are indicated by yellow arrowheads. **(G)** Devoid tubules (from n testes) as percentage of total. **(H)** Sections from 12-month-old testes were analyzed by TUNEL and DAPI staining. TUNEL-positive cells are indicated by yellow arrowheads. **(I)** Percent TUNEL-positive cells per tubule (n indicated by genotype to the right). Statistical significance (*p < 0.05) was determined by one-way ANOVA with correction for multiple comparisons and student’s t-test. Error bars indicate SEM.

To gain insight into spermatogenesis problems that ultimately lead to devoid tubules, we analyzed whether histologically normal tubules contained fewer germ cells or more apoptotic cells at 12 months. Undifferentiated spermatogonia counts, indicated by PLZF staining, did not differ from controls (**Figures S3A, B**), but histologically normal tubules in *Pnma* mutants contained more apoptotic cells, as indicated by the presence of fragmented dsDNA using terminal deoxynucleotidyl transferase dUTP nick end labelling (TUNEL, **Figures 2H, I**). The depleted tubules and germ cell death support the idea that *Pnma1* and *Pnma4* are important for male germ cell development and likely execute their functions prior to spermatid formation.

Like the male mice, female mice lacking *Pnma1* or *Pnma4* were similar to wild type in their reproductive capacity at two months of age, but they birthed smaller litters over time (**Figure 3A, S1E**). This premature loss of fertility correlated with ovarian atrophy, diminished antral follicle counts, and a greater tendency to develop follicular cysts than wild type (**Figures 3B-F, S2B**). Some mutant follicles had lost cumulus granulosa cells, which synthesize steroid hormones and promote oocyte growth, and resembled follicles in mice lacking estrogen receptors (ER) or follicle stimulating hormone (FSH) receptors^22–24^. Other ovarian features did not differ significantly different between mutants and controls with some exceptions (**Figures S4A-E**). Serum progesterone, FSH, and anti-Müllerian hormone (AMH) levels were statistically indistinguishable from controls at 1.5, 3, and 6 months with two exceptions: progesterone was down in *Pnma4^-/-^* at 6 months and AMH was elevated in double mutants at 1.5 months. Oocytes isolated from single and double mutants completed meiosis II at 16 hours at similar frequencies compared to controls, indicating that the reduced fertility in mutant animals is unlikely to be caused by severe meiotic defects (**Figure 3G**).

**Figure 3:**
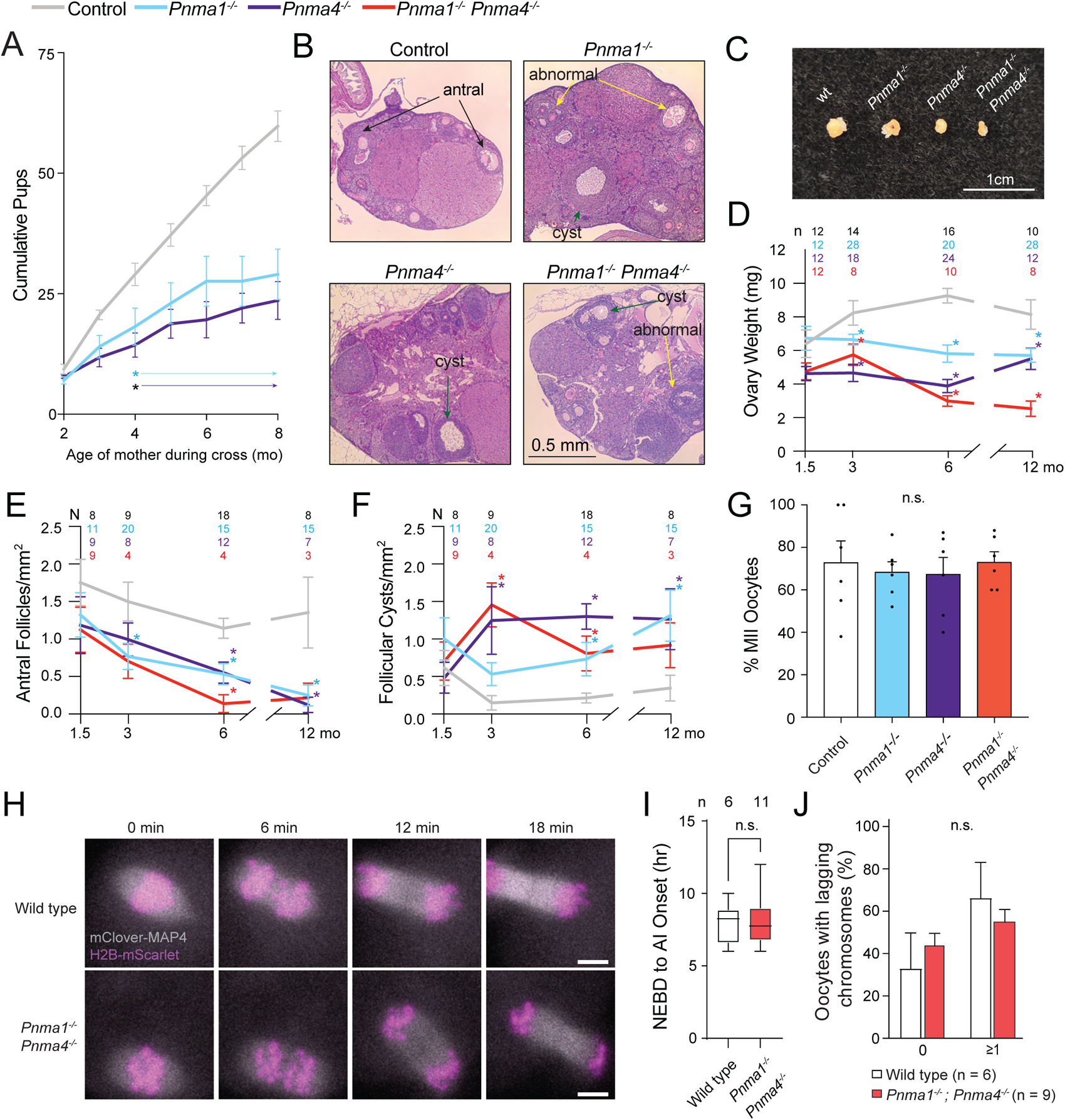
Female mice lacking *Pnma1* or *Pnma4* have age-dependent reproductive defects. **(A)** Wild type female control mice (gray), *Pnma1^-/-^* (cyan), or *Pnma4^-/-^* (purple) were crossed to B6D21/J male fertility tester mice (N = 5 pairs for each genotype-timepoint combination). Pup numbers for each cross are plotted over time. **(B)** PAS-stained ovarian sections from 3-month-old mouse ovaries. Black arrows denote antral follicles, yellow arrows denote abnormal follicles, and green arrows denote follicular cysts. **(C)** Representative images of ovaries from the four genotypes. **(D)** Ovaries (n above graph) from control (pooled wild type and heterozygous, gray), *Pnma1^-/-^* (cyan), *Pnma4^-/-^* (purple), or *Pnma1^-/-^ Pnma4^-/-^* double mutant (red) mice were weighed. **(E)** The number of antral follicles at the indicated ages per mm^2^. The number of individuals (N) for each genotype and age are shown. **(F)** The number of follicular cysts per mm^2^ by age and genotype (N’s in graph). **(G)** Germinal vesicle (GV) oocytes were collected from six-month-old females and meiotically induced. Meiotic progression was analyzed by tubulin immunofluorescence. Percentage of meiosis II oocytes was recorded. **(H)** GV oocytes were collected from seven-month-old wild-type and double mutant mouse ovaries. We injected mRNA encoding *mClover-MAP4* (microtubule-binding protein) and *H2B-mScarlet* (histone) and imaged oocytes live for ∼18 hours, recording **(I)** the time from nuclear envelope breakdown (NEBD) to anaphase I and **(J)** lagging chromosome percentages. Statistical significance (*p < 0.05) was determined by determined by one-way ANOVA with correction for multiple comparisons and student’s t-test. Error bars indicate SEM.

To determine whether mutant oocytes that do complete both meiotic divisions have subtler meiotic defects, we used high-resolution live imaging to analyze chromosome segregation during oocyte maturation. Oocytes from double mutants at two and seven months did not differ from controls in meiotic chromosome segregation, alignment on the metaphase I and metaphase II spindles, or progression timing (**Figures 3H-J, S5A-G**), suggesting that mutant oocytes that do progress are meiotically competent. To assess the mouse ovarian cell types where *Pnma* genes may act, we analyzed the expression of *Pnma1-5* in single cells at young (4.5 months), peri-estropause (10.5 months), and post-estropause (15.5 months) stages (**Figure S6**). *Pnma1* and *Pnma4* showed the highest expression levels throughout the ovary compared to other *PNMA* genes (except *Pnma5* specifically in peri-estropause oocytes, **Figure S6**, orange dots). *Pnma1* and *Pnma4* are primarily co-expressed (by % positive cells) in granulosa, stroma, and theca cells. Orthogonal analysis of *Pnma1* and *Pnma4* expression from a published data set^25^ corroborated that these genes are expressed in subpopulations of granulosa and stroma cells (**Figures S7A, B**). In granulosa, stroma, and theca, we observed a ∼10-fold decline in *Pnma1*-positive cells at post-estropause (**Figure S6**, gray dots). *Pnma4* expression did not change significantly in the same time frame. Together, these data support the idea that these genes are synthesized in subpopulations of ovarian cells that support oocyte maturation and viability.

### *Pnma1* and *Pnma4* mutants acquire abdominal obesity but appear behaviorally normal

In humans and mice, reproductive decline that occurs naturally with age commonly correlates with changes in body composition^26,27^. As male and female *Pnma* mutants aged, we noticed that they rapidly gained abdominal subcutaneous and visceral fat, corresponding to the time course of their dwindling fertility. At 1.5 months, the knockouts were statistically indistinguishable from controls, but by 3 months, *Pnma1^-/-^* and double mutants were 20% heavier than controls (**Figures 4A**, **B**). By 6 months, all mutants were significantly heavier than controls. Mutant mice did not consume more food compared to controls (**Figure 4C**), so the weight gain likely reflects altered metabolism. A similar phenotype is observed post-ovariectomy^28^, in mice lacking sex hormone receptors^22–24^, and in castrated males^29^. In human and rodent males, increased adiposity arises due to testosterone deficiency^30^.

**Figure 4:**
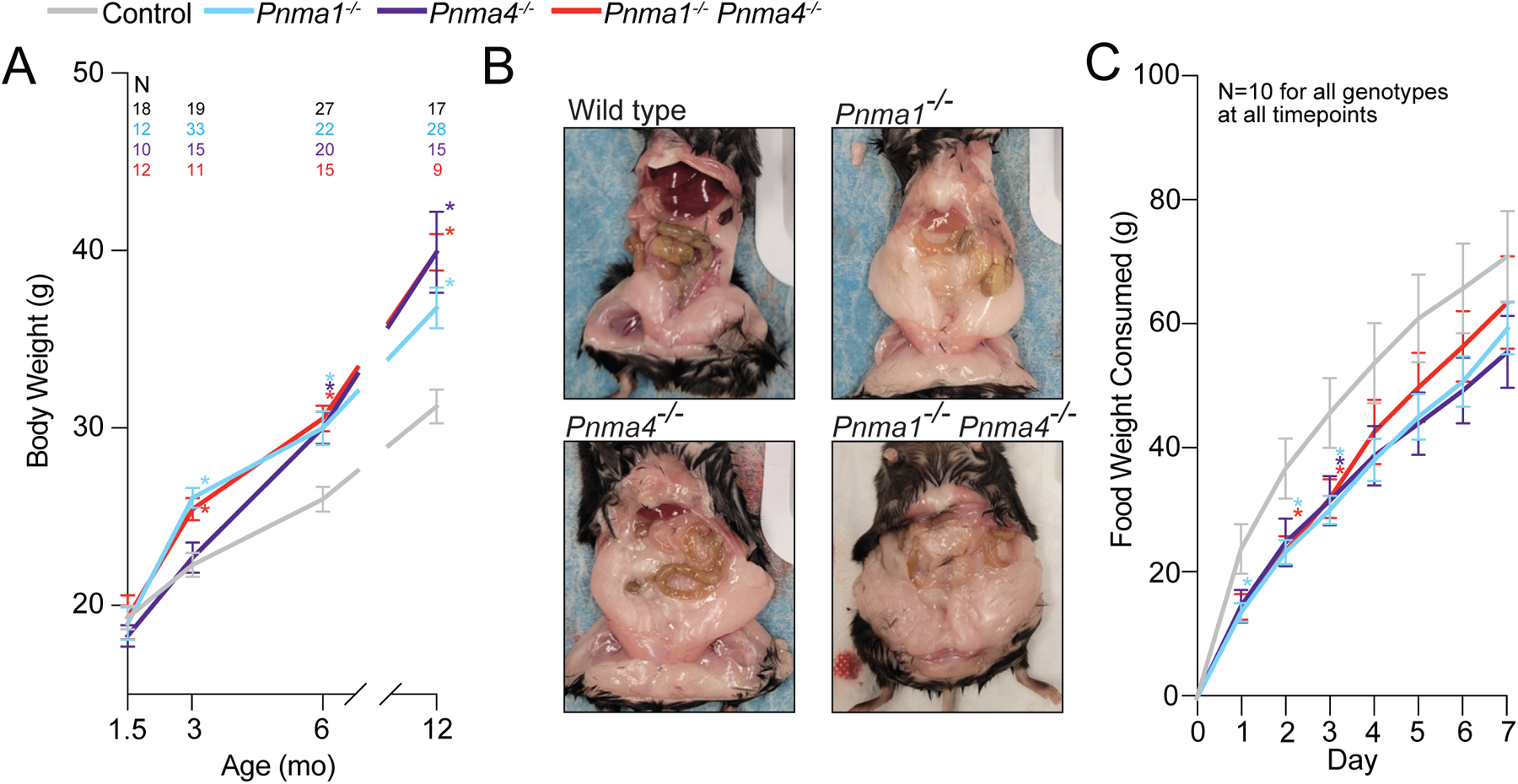
*Pnma1* and *Pnma4* mutants gain abdominal fat with age. **(A)** Body weights (N above graph) of control (pooled wild type and heterozygote, gray), *Pnma1^-/-^* (cyan), *Pnma4^-/-^* (purple), or *Pnma1^-/-^ Pnma4^-/-^* double mutant (red) at the indicated ages. **(B)** Representative images of abdominal fat at six months in each genotype. **(C)** Food intake of 3-month wild type, *Pnma1^-/-^*, *Pnma4^-/-^*, or *Pnma1^-/-^ Pnma4^-/-^* double mutant mice (N = 10 mice) was monitored daily for 7 days. Statistical significance (*p < 0.05) was determined by one-way ANOVA with correction for multiple comparisons. Error bars indicate SEM.

Like other *PNMA* genes, both *PNMA1* and *PNMA4* are expressed in the brain^31^. We therefore tested basic neurological functions in six-month-old male *Pnma1^-/-^*, *Pnma4^-/-^*, and double mutants compared to wild type. The Y-maze spontaneous alternation assay revealed no deficits in short-term or spatial memory (**Figure S8A**). Mutant mice were largely indistinct from controls regarding long-term memory, as assessed by the amount of post-cue freezing behavior after fear conditioning (**Figure S8B**). In the open field test, mutants did not differ significantly from controls in time spent in the center, distance traveled, and vertical counts (**Figure S8C-E**). We observed no difference in muscular strength or coordination between single and double mutants in the inverted hang or grip strength tests (**Figure S8F, G**). Although these results revealed no obvious behavioral abnormalities in *Pnma1^-/-^* and *Pnma4^-/-^* male mice, it is possible that female mutants or mice at other ages could display differences in cognitive function.

### PNMA1 and PNMA4 form capsid-like structures in gonadal tissue

Gag family proteins self-assemble into capsid-like structures that are critical for packaging and delivering molecular cargo. One prominent example is ARC, which has been independently domesticated in mammals and *Drosophila* to encapsulate and traffic mRNA across neuronal synapses^32,33^. To determine whether PNMA1 and/or PNMA4 proteins have retained the ability to self-assemble into capsids, we expressed and purified His-tagged recombinant (mouse) PNMA1 and PNMA4 from *E. coli* which we analyzed by size exclusion chromatography (**Figures S9A, B**). *Metaviridae* capsids are amenable to isolation by size-based fractionation because they are large (yeast Ty3 capsids contain 540 gag protein subunits and are ∼ 20 megadaltons^34^). We visualized the high molecular weight fractions by negative stain transmission electron microscopy (TEM). Non-uniform capsid-like assemblies are evident in micrographs of both PNMA1 and PNMA4 (16-21 nm diameter for PNMA1 and 36-51 nm for PNMA4, **Figures 5A, B**). Based on their size, these results suggest that PNMA1 forms capsids similar to those formed by PNMA2^35^ , while PNMA4 capsids are similar to those formed by LTR retroelements^34^.

**Figure 5:**
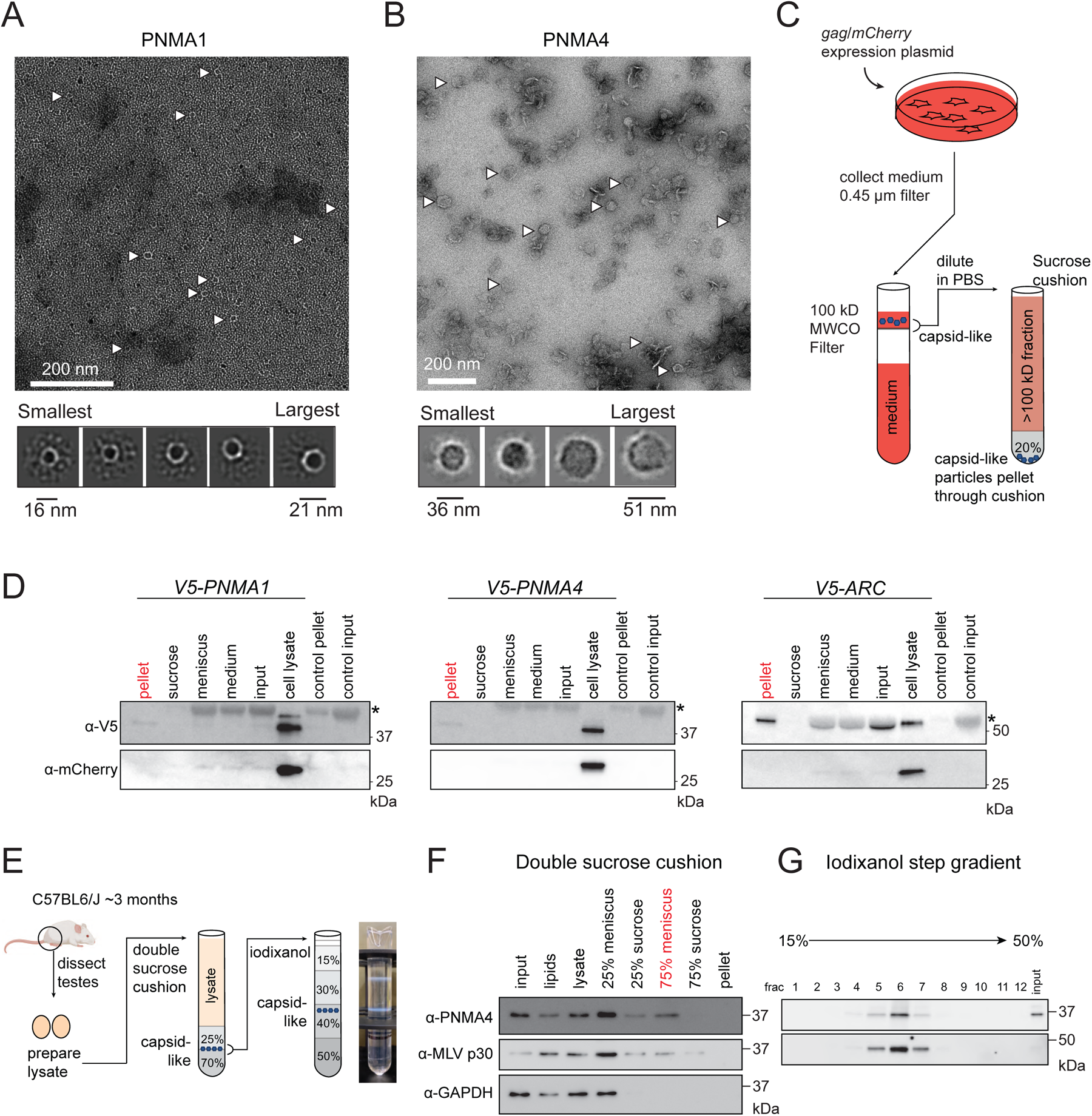
PNMA proteins form capsid-like structures that can exit human cells. **(A, B)** Top panels depict TEM micrographs of negatively stained recombinant PNMA1 (A) and PNMA4 (B) protein. Bottom panels depict 2D class averages from cryoSPARC. **(C)** Experimental setup: *3V5*-tagged *PNMA1*, *PNMA4*, and *ARC* (control exported capsid) expression plasmids were transfected into HEK-293T cells. *mCherry* (control non-capsid) was co-expressed from the transfected plasmid. Culture medium was collected, spun, and filtered to remove cells and debris. Filtrate was fractionated over a 20% sucrose cushion to enrich for capsids, which fractionate as large particles at the bottom of the cushion (pellet). Sucrose, meniscus, spun media, and pellet fractions were collected. Unfractionated media input and cell lysate samples were also collected. **(D)** PNMA, PNMA4, ARC (anti-V5), and mCherry protein levels by immunoblot. Untransfected controls (control input) are at the far right of each blot. Asterisks (*) mark the cross-reacting BSA band (abundant in growth medium). Note the presence of BSA in the untransfected controls and absence of BSA in pellet/lysate. **(E)** Experimental setup: Lysate was prepared from ten testes from wild type C57BL6/J mice were collected at 3 months. Lysate was fractionated by velocity step gradient (38,000 rpm, 3 hours) over a double sucrose cushion (25% and 70%). Capsid-like structures migrate to the interface between 25%-70% sucrose (70% meniscus). The 70% meniscus was further fractionated by isopycnic (equilibrium) centrifugation on an iodixanol step gradient; the photograph shows the centrifuge tube after this step. **(F-G)** PNMA4, MLV p30 gag, and GAPDH protein levels were determined by immunoblot in each fraction at the end of the double sucrose fractionation and **(G)** iodixanol fractionation.

To assess capsid formation in mammalian cells, we expressed human *PNMA1*, *PNMA4* and *ARC* (positive control) in HEK293T cells with *mCherry* (non-capsid control, driven by a separate promoter). After 24 hours, we harvested cells and fractionated cell lysate by velocity gradient ultracentrifugation. Because of their size and density, capsids migrate through the gradient, away from the major protein content of the cell. Some PNMA1 and PNMA4 appeared in denser factions (**Figure S10**). To determine if these large assemblies have the capacity to exit cells, we expressed *PNMA1*, *PNMA4* and *ARC* for 24 hours in human cells, then collected the growth medium and removed the cells and any debris by filtration. We concentrated the medium using a 100 kDa molecular weight cutoff (MWCO) filter, discarded the < 100 kDa flow-through, fractionated the concentrate by sedimentation over a 20% sucrose cushion, and analyzed the fractions by immunoblot (**Figures 5C, D**). Like ARC, some PNMA1 and PNMA4 appeared in the pellet fraction (**Figure 5D**), supporting the idea that at least some PNMA proteins exit human cells in a capsid-like form.

To assess whether PNMA proteins form capsid structures *in situ*, we purified PNMA4 capsids from wild-type mouse testes using a procedure that was developed for the purification of intact VLPs from plant tissue^36^. We focused on PNMA4 because we found its detectability in testes by (polyclonal) immunoblot more reliable than PNMA1. To separate PNMA4 capsids from monomers and oligomers, we fractionated testis lysate over two sucrose cushions of 25% and 70% by ultracentrifugation (**Figure 5E**). Because of their size and density, capsid particles migrate to the interface between the 25% and 70% sucrose layers and separate from monomeric proteins, which accumulate at the interface between the lysate and 25% cushion^36^. We observed enrichment of PNMA4 at the 25%-70% interface compared to a control monomeric protein (GAPDH) and co-enrichment with p30 gag from murine leukemia virus (MLV), an enveloped capsid positive control (**Figure 5F**). To separate capsids from other large structures such as ribosomes, we collected and re-fractionated the 25%-70% meniscus fraction by isopycnic (equilibrium) ultracentrifugation on an iodixanol (Optiprep) density step gradient. PNMA4 protein isolated from testis tissue co-fractionated (as large particles) with MLV p30 capsid protein (**Figure 5G**). The *in situ* biochemical properties of PNMA4 are consistent with capsid formation, which may mediate the protein’s pro-fertility function.

### *PNMA1* and *PNMA4* expression in human ovaries declines with age

While previous studies of mouse male gonads indicated that both *PNMA1* and *PNMA4* are expressed at their highest levels in spermatocytes^37^, analysis of mouse female gonads (here and published^25^) indicated that both genes are primarily expressed in follicular cells. We therefore sought to determine which and to what degree human female gonadal cells express *PNMA*-family genes. We analyzed *PNMA1-5* expression by single-nuclei RNAseq in human ovaries donated from four young individuals in their 20s, four older individuals in their late 40s to early 50s, and in published human oocyte expression data^38,39^. As in mice, *PNMA1* and *PNMA4* showed the highest expression levels throughout the ovary (**Figures 6A, B, S11**). Oocytes expressed the highest levels of *PNMA1* and *PNMA4*. The somatic ovarian cell type that expressed the highest levels of either gene was granulosa cells: in young adult ovaries, 8.5% of granulosa cells expressed *PNMA1* and 5.1% expressed *PNMA4*, possibly indicating specific granulosa subpopulations (**Figure 6B**). Comparing younger and older ovaries, *PNMA1* expression was significantly higher in young granulosa, stroma, and blood endothelium cells, while *PNMA4* was significantly higher in young smooth muscle and blood endothelium cells, signifying age-related decline in expression of these genes.

**Figure 6:**
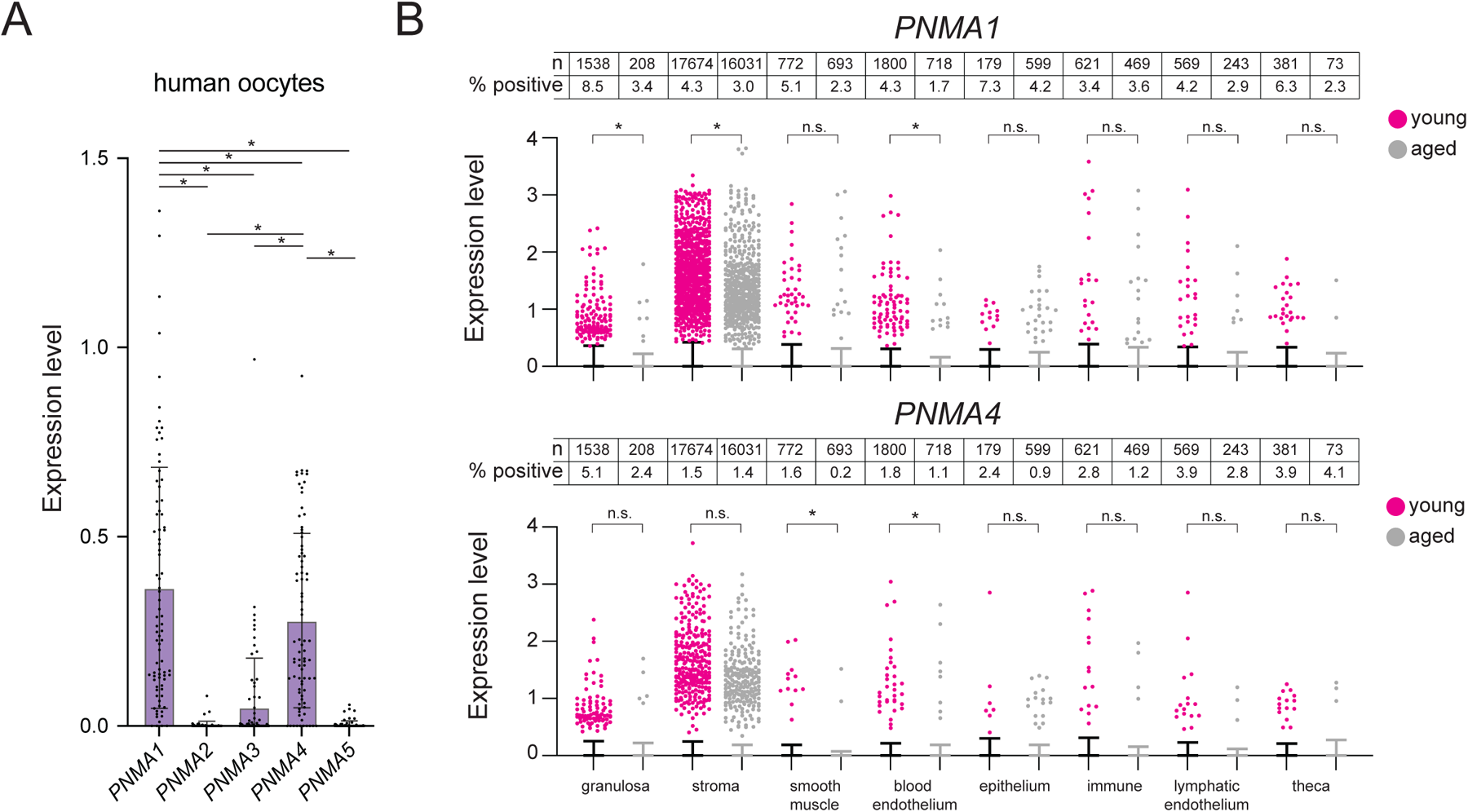
*PNMA1* and *PNMA4* are expressed in human ovaries. **(A)** Analysis of uniquely mapping single-cell RNAseq reads for *PNMA1-5* loci expressed in MII human oocytes (data from Yuan et al., 2021)^38^. **(B)** Human ovaries from young (23-29 years of age, N = 4) and older (49-54 years of age, N = 4) donors were analyzed by single-nuclei RNAseq. Uniquely mapping reads for *PNMA1* and *PNMA4* loci were assigned to ovarian tissue types based on clustering analysis. Note that the majority of values = 0 and lie underneath the x axis (error bars represent SEM). Total n of each cell type and percentage of positive cells is indicated above the plots. Statistical significance (*p< 0.05) was determined by Mann-Whitney test with Welch’s correction.

In mouse spermatocytes and human fetal ovaries, the 3’ untranslated regions (UTRs) of *PNMA1* and *PNMA4* transcripts are bound by DAZL^40,41^, which can activate or repress mRNA translation^42^. To test whether the PNMA-DAZL interaction could be of regulatory significance, we expressed human *PNMA1*, *PNMA4*, or *SMC1b* (control DAZL target) with or without DAZL in human cells (HEK293T, **Figure S12A**). DAZL augmented *PNMA1* protein levels and repressed *PNMA4* without altering mRNA levels (**Figure S12B, C**).

### Human variation at the *PNMA1* and *PNMA4* loci is associated with reproductive defects

Based on the reduction of fertility in mice lacking *Pnma1*, *Pnma4*, or both, we asked whether subfertility or other reproductive phenotypes in humans are causally linked to genetic variation in the *PNMA1* and/or *PNMA4* loci (**Table S1**). We used the variant-to-gene (V2G) and locus-to-gene (L2G) pipelines available from Open Targets Genetics, with a standard p-value cutoff of 5x10^-8^, to identify the causal gene from a trait-associated variant or locus identified by GWAS^43,44^. Briefly, from a trait-associated variant (*i.e.*, the polymorphism corresponding to a GWAS peak), these pipelines establish the likely causal gene by weighted analysis of independent published quantitative trait loci experiments (eQTLs, pQTLs and sQTLs), chromatin interaction experiments, *in silico* functional predictions, and distances between a variant and transcription start sites. We found six variants attributed to *PNMA1* (from nine studies and three publications^45–47^) associated with altered testosterone levels and levels of sex hormone binding globulin (SHBG, levels correlate to testosterone). Two variants attributed to *PNMA4* are associated with altered age of menarche^48^ and male puberty timing^49^. We also identified two additional variants associated with elevated body fat (thirteen studies, three publications^50–52^) that are attributed to *PNMA4*. These results support the notion that the alterations in reproductive capacity we observed in mouse *Pnma1* and *Pnma4* knockouts are relevant to humans.

## DISCUSSION

In general, retroelement gag capsids facilitate endogenous and parasitic intercellular communication. Like many other tissues, gonads rely on endocrine, paracrine, and autocrine signaling to support gamete development, and their cells employ a variety of means to communicate with each other. For example, testosterone, produced largely in Leydig cells, diffuses into the testicular tubules where it binds to androgen receptors on Sertoli cells which then drive the production of factors that support spermatogenesis^53,54^. Progesterone, produced by follicular granulosa cells and corpora lutea, is critical for successful oocyte maturation and ovulation^55,56^. Communication among germ cells also takes place through intercellular bridges that connect the cytoplasm of adjacent cells into a syncytium; males that cannot form these bridges (such as *Tex14* knockouts) are sterile^57^ (these structures are important, but not essential, for female fertility^58,59^). These germ cell bridges/syncytia have been proposed to foster dosage compensation in haploid cells, coordination of meiotic entry, and sharing of signals for synchronous cell divisions within seminiferous tubules^57,58,60^. In the ovary, oocytes are surrounded by cumulus/coronal granulosa cells that extend long processes termed transzonal projections through the zona pellucida^61–63^. These projections release vesicles that are thought to contain RNA and have been proposed to enable sharing of RNA between transcriptionally active granulosa cells and the silent oocyte^64,65^. Conversely, oocytes release growth factors that regulate proliferation of granulosa cells. Emerging evidence supports the notion that communication between follicular cells and oocytes is important for maintaining germ cell fitness: aged oocytes transplanted into young follicles become rejuvenated, improving in mitochondrial function and the fidelity of meiotic chromosome segregation, among other measures^66^. Although PNMA1 and PNMA4 are not *essential* for intercellular communication (the double mutant would be sterile), we propose that PNMA1 and PNMA4 capsids could promote the transmission of cytoplasmic signals, whether through bridges, transzonal projections, and/or by transport across plasma membranes (**Figure 7**).

**Figure 7:**
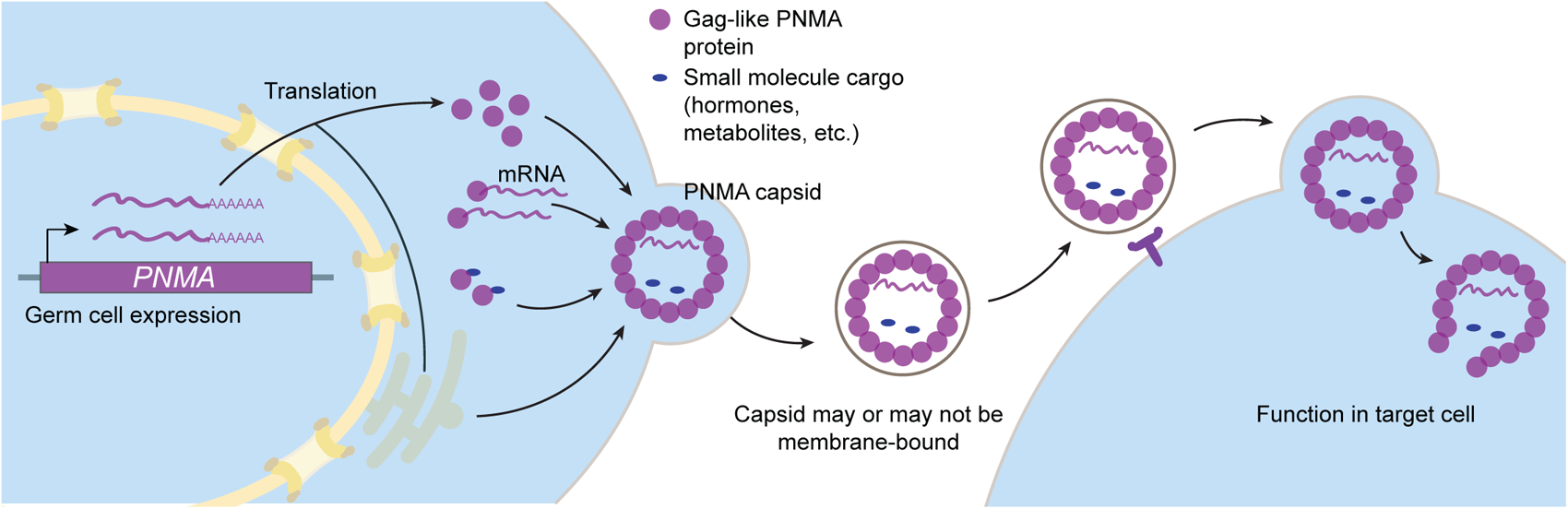
Model for PNMA1 and PNMA4 function.

The word capsid derives from the Latin word for “case,” and capsids form an elegant means to carry molecular messages. The question we still need to answer is, do the capsids formed by PNMA1 and PNMA4 carry cargo, and if so, what is it? This cargo could conceivably be nucleic acids such as mRNA or tRNA, hormones such as testosterone or FSH, or a metabolite—but whatever the cargo, it likely has important functions for fertility.

The literature provides a few clues as to the likely purpose of the possible cargo. *PNMA4* has been implicated in apoptosis in the context of mis-expression in cancer cells^67–69^. PNMA4 interacts with the pro-apoptotic protein Bax and the pro-survival proteins Bcl-2 and Bcl-X(L) and can mediate caspase-dependent apoptosis when overexpressed^70^. Given that 12-month-old *Pnma4*^-/-^ testes exhibit greater numbers of TUNEL-positive cells, *PNMA4* likely promotes survival in gonadal tissue. In mutant males, we observed an age-dependent intra-organ heterogeneity where some testicular tubules were devoid of sperm and sperm precursor cells while others appeared wild type. Other than increased cell death in aged tubules, we did not observe any clear defects in the ‘normal’-appearing seminiferous tubules of mutant animals. Because mutant tubules are either devoid or full, without an obvious intermediate state, we speculate that devoid tubules have experienced a failure to initiate a wave of spermatogenesis and, once a failure has occurred, that tubule cannot recover.

Female mice lacking *Pnma1* and/or *Pnma4* exhibit a wide range of ovarian abnormalities. *Bmp6^-/-^, Bmp15^-/-^, Smad2/3^cko^,* and *Smad3^cko^* female mice exhibit age-dependent subfertility with ovarian histology reminiscent of *Pnma* knockouts^71–74^, suggesting that the *Pnma1* and *Pnma4* mutant phenotypes could, in part, be due to downregulated TGF-beta signaling. In mutant females, the oocytes that are produced appear healthy and undergo meiosis. Similarly, while sperm count declines with age in mutant males, the sperm within the epididymis appear normal. These data are consistent with a model in which the function(s) of PNMA1 and PNMA4 are enacted prior to gamete release from the ovary/testis though we cannot rule out defects in later processes such as implantation. Elucidating their functions will likely require identifying the cargo that is being transported.

Developing yeast gametes thwart proliferation of *Ty3*, a parasitic ancestor of *PNMA* genes, by using amyloid-like condensates of the RNA-binding protein Rim4, which binds retrotransposon mRNA and inhibits its translation^10^. Mammalian analogs of Rim4 include DAZ, DAZL (DAZ-like), and PUM1/2 (Pumilio). Although higher-order assemblies of RNA-binding proteins behave somewhat differently in yeast and mammalian germ cells, DAZ, DAZL, and PUM1/2 are like Rim4 in that they regulate translation in developing gametes, affect gene expression post-transcriptionally by binding the UTRs of their target genes, and form condensates *in vivo*^75–78^. Our analysis of DAZL-mediated regulation of *PNMA* genes supports the idea that, from yeast to humans, RNA-binding proteins with the propensity to form condensates can regulate the translation of retroelement-derived mRNA. We propose that as *Metaviridae* elements gradually evolved beneficial functions and lost proliferative capacity in host meiotic cells, the analogs of condensates acting as a defense mechanism (Rim4) were repurposed in mammals as regulators (DAZL). Could DAZL have maintained a role to protect sperm and eggs from the assault of expressed *Metaviridae* retrotransposons? Several human endogenous retroviruses (hERVs) are robustly expressed in gonads^79,80^ and these elements do not appear to be rapidly proliferating, suggesting that the hERV lifecycle is somehow blocked after mRNA transcription. The possibility for condensate-forming RNA-binding proteins to post-transcriptionally regulate hERVs has precedent: in neurons, the neurodegeneration-associated protein TDP-43 binds to hERV-K mRNA and regulates its protein synthesis^81^.

It has been noted that PNMA capsids should be suitable as delivery vehicles for therapeutic agents such as RNA vaccine delivery and gene therapy^18,19,32,82^; such therapies involve bioengineering capsid proteins for the delivery of designer cargo, such as mRNA. The beauty of PNMA1 and PNMA4 is that they would require no engineering to function as therapeutic agents for subfertility because they could naturally harbor pro-fertility cargo. Therapeutic development would focus on PNMA4 because, unlike PNMA1, it is not associated with autoimmunity^83^. Before a therapy for subfertility can be developed, however, we must optimize methods to purify PNMA4 capsids from gonadal tissue, understand precisely what cargos they carry, and assess their therapeutic efficacy in mouse models of subfertility.

## SUPPLEMENTAL FIGURE LEGENDS

**Figure S1:**
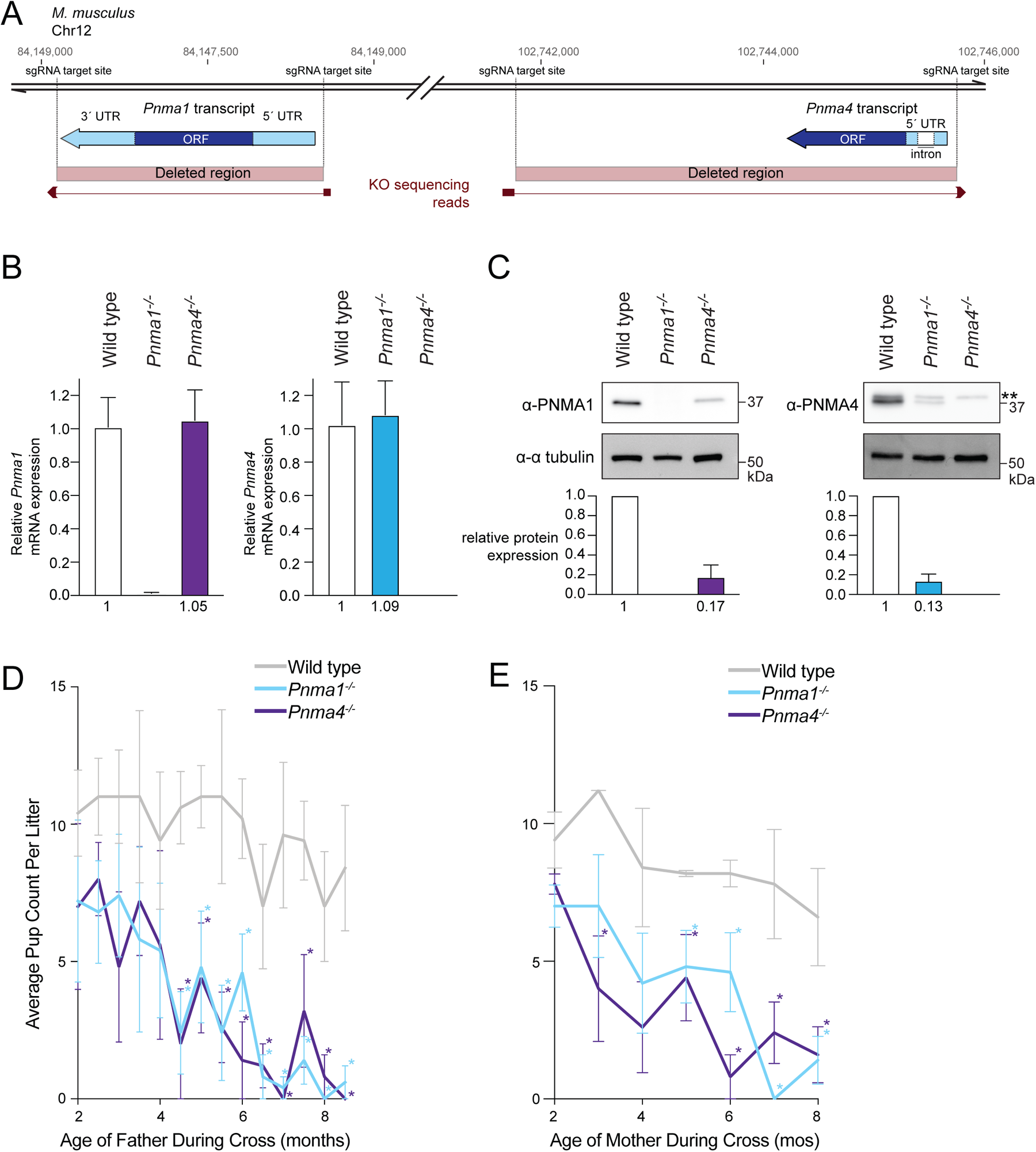
Diagram of *Pnma1* and *Pnma4* loci on *M. musculus* chromosome 12. **(A)** *Pnma1* and *Pnma4* deletions were generated by CRISPR/Cas9 genome editing (sgRNA target sites shown in dashed lines). Successful deletion was assessed by PCR/sequencing. Shown are chromosomal coordinates, annotated transcripts (CDS in navy, UTRs in cyan, and introns in white), deleted regions, and sequencing reads (below, maroon) confirming deletion of *Pnma1* and *Pnma4*. **(B, C)** Analysis of mRNA and protein produced in the *Pnma1^-/-^* and *Pnma4^-/-^* mutants. A testis was dissected from 3-month wild type (C57BL/6J), *Pnma1^-/-^*, and *Pnma4^-/-^* mice. Lysate was prepared and split for mRNA and protein analysis. **(B)** Analysis of *Pnma1* and *Pnma4* mRNA levels by qPCR (n = 3 replicates). **(C)** Analysis of PNMA1, PNMA4, and α-tubulin (loading) protein levels by immunoblot. Shown below is a quantification of the ratio of PNMA1 and PNMA4 in knockouts vs. wild type (set to 1) from n = 3 biological replicates. ** denotes presence of an unfortunate cross-reacting band present in all lanes of the α-PNMA4 immunoblot. Error bars indicate SEM. **(D)** Male wild type mice (gray), *Pnma1^-/-^* (cyan), or *Pnma4^-/-^* (purple) were crossed biweekly to CF-1 female fertility tester mice (N = 5 pairs for each genotype-timepoint combination). **(E)** Female wild type mice (gray), *Pnma1^-/-^* (cyan), or *Pnma4^-/-^* (purple) were crossed to B6D21/J male fertility tester mice (N = 5 pairs for each genotype-timepoint combination). Pup numbers for each cross were recorded. This is the same experiment used to generate the data in Figures 2A and 3A, but here average pup count per litter is plotted. Statistical significance (*p < 0.05) was determined by student’s t-test. Error bars indicate SEM.

**Figure S2:**
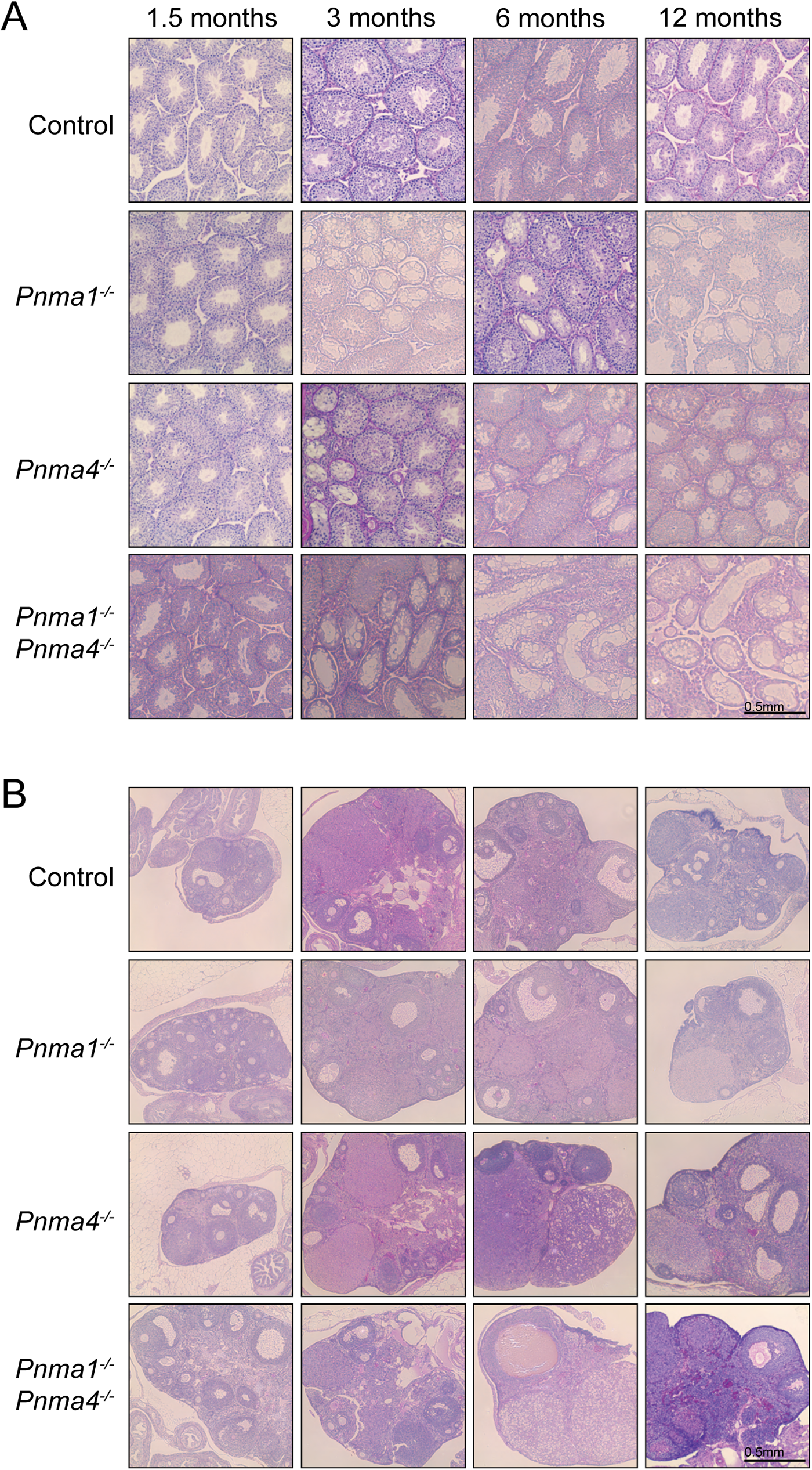
Defective testicular and ovarian morphology in *Pnma1* and *Pnma4* mutants. **(A)** Representative PAS-stained testis sections by age. Quantified in Figure 2. **(B)** Representative PAS-stained ovary sections by age. Quantified in Figure 3 and S4.

**Figure S3:**
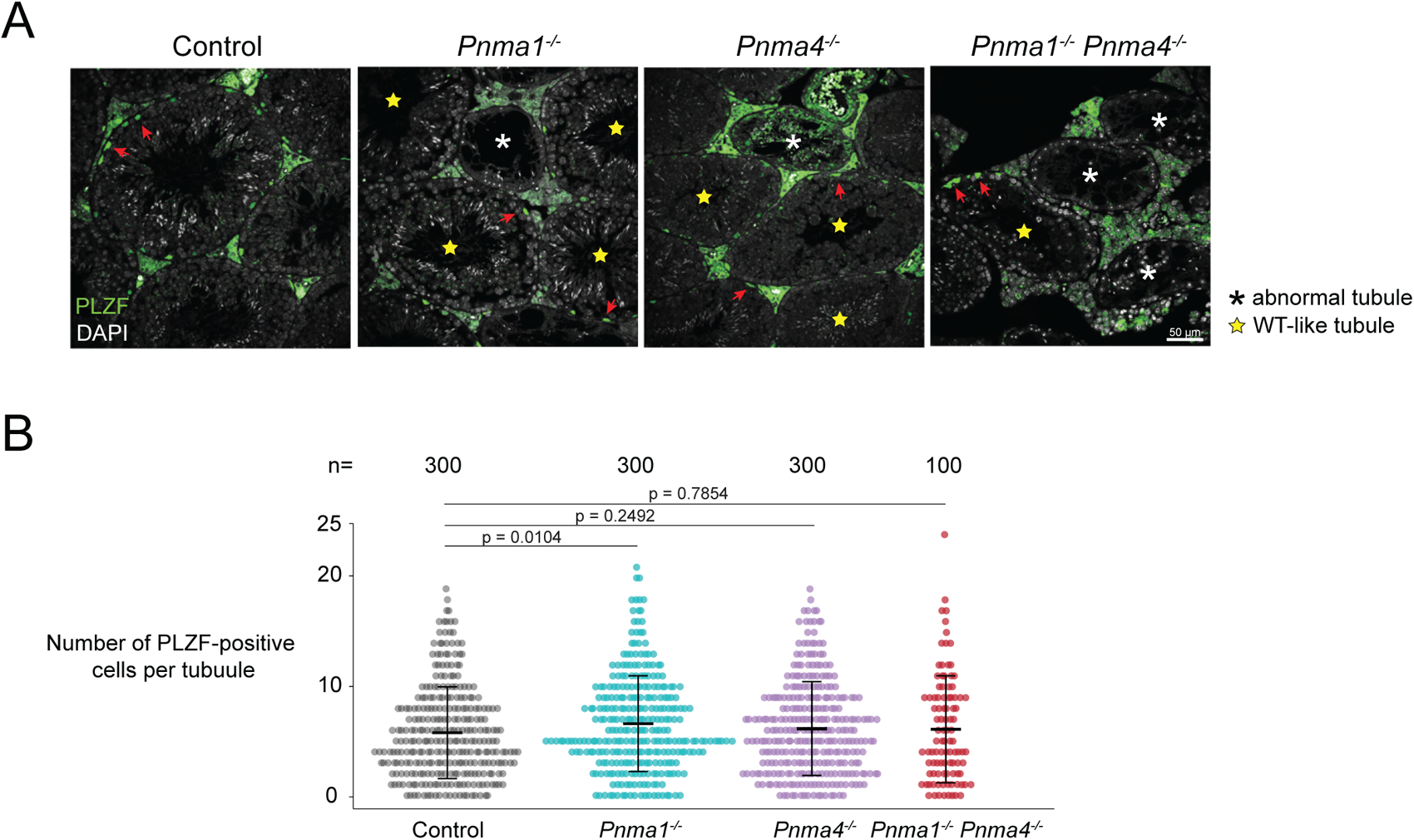
Analysis of undifferentiated spermatogonial cells by PLZF staining. **(A, B)** Testes were fixed, paraffin embedded, and sectioned. Sections from 12-month samples were analyzed by TUNEL and DAPI staining. **(A)** Images and **(B)** quantifications (from n wild type-like tubules) of PLZF-positive cells (red arrows) are shown. Statistical significance (*p < 0.05) was determined by Mann-Whitney test. Error bars indicate SD.

**Figure S4:**
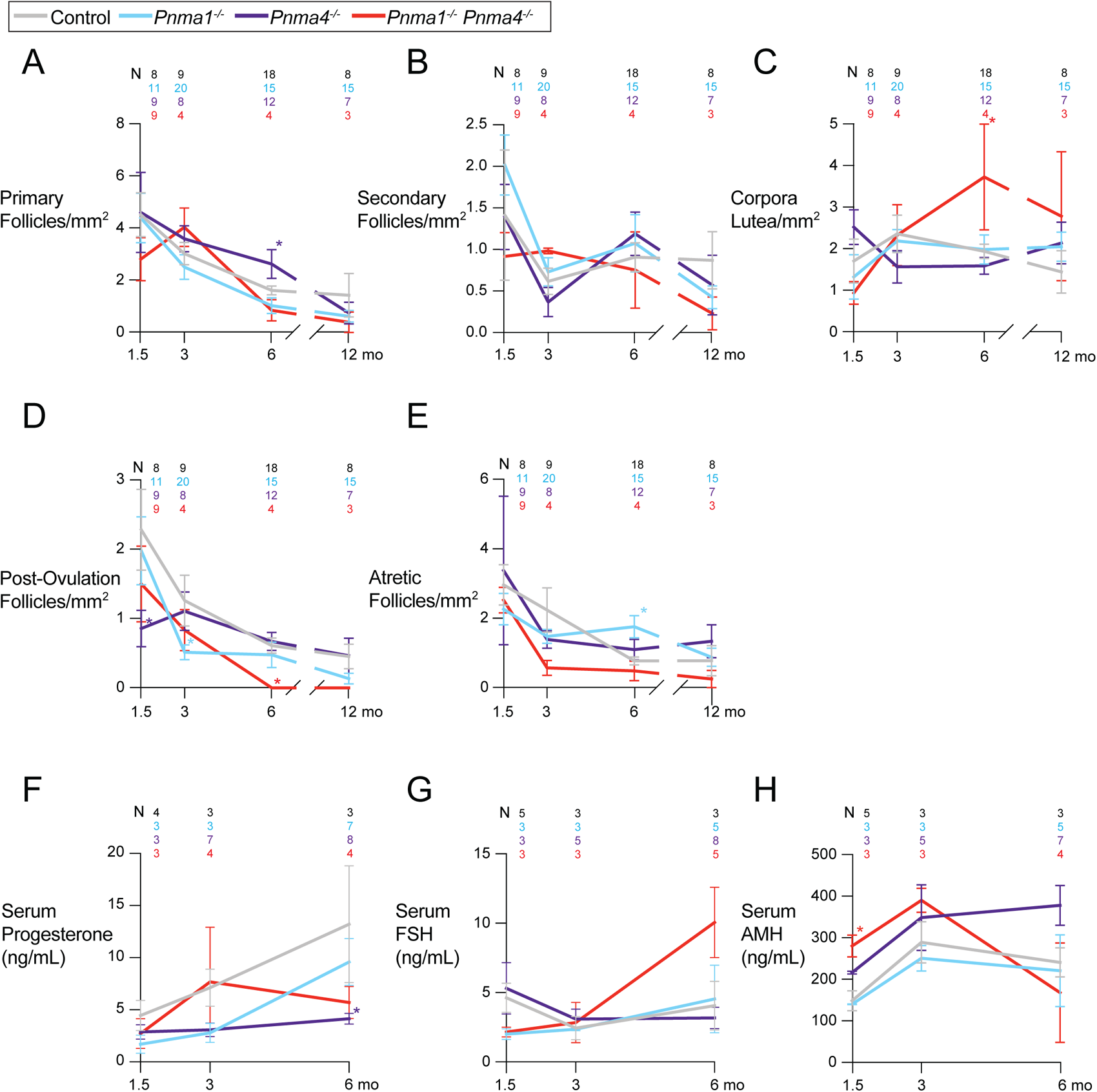
Several ovarian features are not dramatically altered in *Pnma1* and *Pnma4* mutants. **(A-E)** Ovaries from control (pooled wild type and heterozygote, gray), *Pnma1^-/-^* (cyan), *Pnma4^-/-^*(purple), or *Pnma1^-/-^*; *Pnma4^-/-^* double mutant (red) mice were fixed, embedded, and PAS-stained. The following features were quantified per unit area (mm^2^) at the indicated times: **(A)** primary follicles, **(B)** secondary follicles, **(C)** corpora lutea, **(D)** post-ovulation follicles, and **(E)** atretic follicles. **(F-H)** Serum levels of **(F)** progesterone, **(G)** FSH, and **(H)** AMH from females was measured at the indicated ages. Statistical significance (*p < 0.05) was determined by determined by one-way ANOVA with correction for multiple comparisons. Error bars indicate SEM.

**Figure S5:**
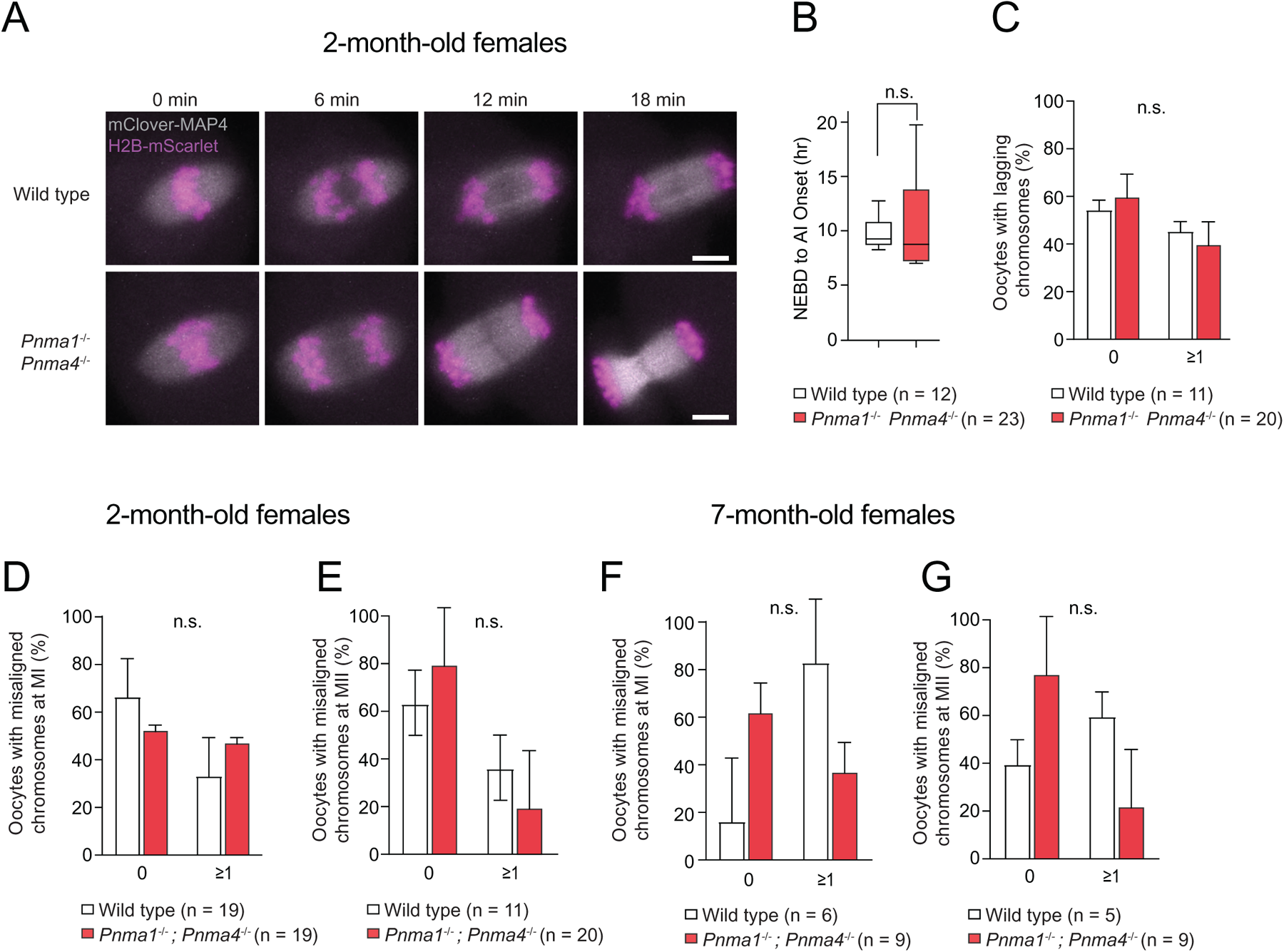
Oocytes that enter meiosis are largely unaffected in *Pnma1* and *Pnma4* mutants. GV oocytes were collected from ovaries of two-month-old wild-type and *Pnma1^-/-^*; *Pnma4^-/-^* mice. We then injected mRNA encoding *mClover-MAP4* (microtubule-binding protein) and *H2B-mScarlet* (histone) and imaged oocytes live for ∼18 hours. **(A)** Representative images of the first meiotic division from the metaphase I (MI) plate (t = 0 min) to anaphase I (AI, t = 20 min). Misaligned chromosomes were counted at t = 0 min and lagging chromosomes at t = 12 min. **(B)** Time in hours (hr) between nuclear envelop breakdown to AI onset. **(C)** Percent of oocytes with no lagging chromosomes or more than one lagging chromosome during AI. **(D)** Percent of oocytes with no misaligned chromosomes or more than one misaligned chromosome in MI plate. **(E)** Percent of oocytes with no misaligned chromosomes or more than one misaligned chromosome at Metaphase II (MII) plate, 4 hours after meiosis I completion. **(F, G)** Continued data from Figure 3. As in A-E, but GV oocytes were collected at 7 months. (F) Percent of oocytes with no misaligned chromosomes or more than one misaligned chromosome in MI plate. (G) Percent of oocytes with no misaligned chromosomes or more than one misaligned chromosome at MII plate, 4 hours after meiosis I completion.

**Figure S6:**
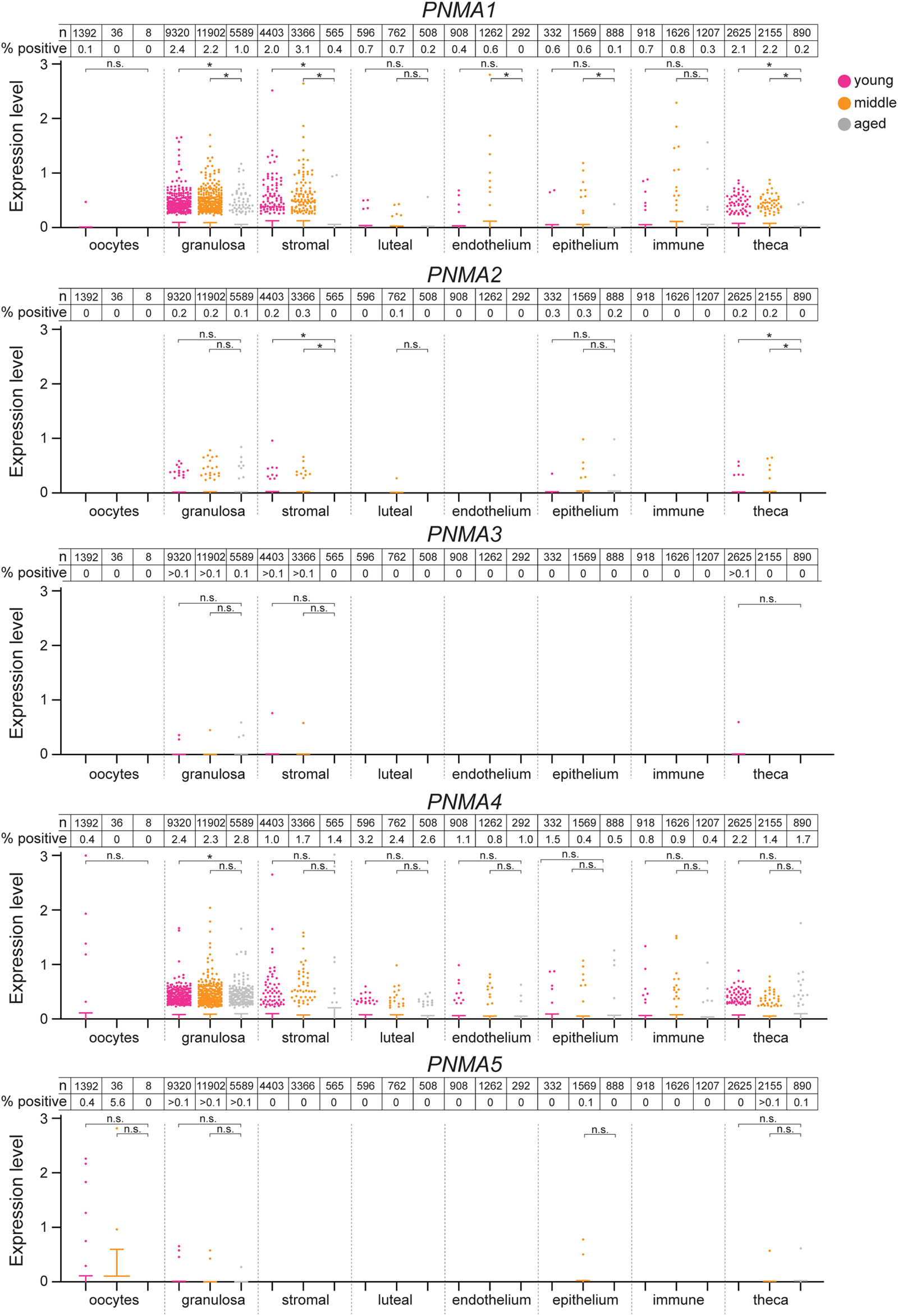
Single-cell analysis of *Pnma1* and *Pnma4* expression in aging mouse ovaries. Ovaries from young (4.5 month, magenta, N = 5), peri-estropause (10.5 month, orange, N = 7), and post-estropause (15.5 month, gray, N = 5) wild type C57BL/6J mice were analyzed by single-cell RNAseq. Uniquely mapping reads for *Pnma1-5* loci were assigned to ovarian tissue types based on clustering analysis. Note that the majority of values = 0 and lie underneath the x axis (error bars represent SEM). Total n of each cell type and percentage of positive cells is indicated above the plots. Statistical significance (*p < 0.05) was determined by Mann-Whitney test with Welch’s correction. Comparisons were not run between conditions with no positive cells.

**Figure S7:**
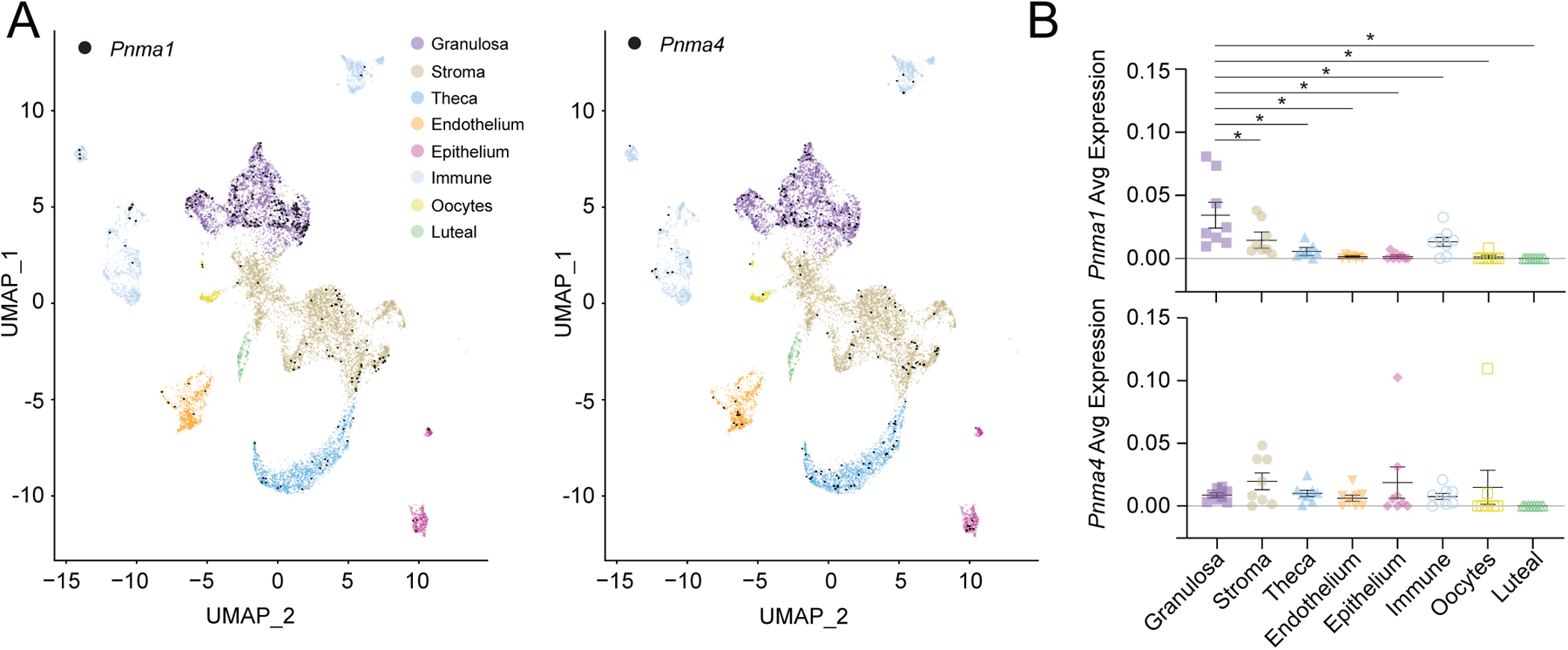
*Pnma1* and *Pnma4* are expressed in subpopulations of ovarian cells. **(A, B)** single cell RNA-seq was performed in n = 8 ovaries from adult C57BL/6J mice. **(A)** UMAP plot showing distinct ovarian cell populations represented by different colors with *Pnma1* and *Pnma4*-positive cells denoted in black. **(B)** Average expression of *Pnma1* and *Pnma4* by sample across different ovarian cell populations.

**Figure S8:**
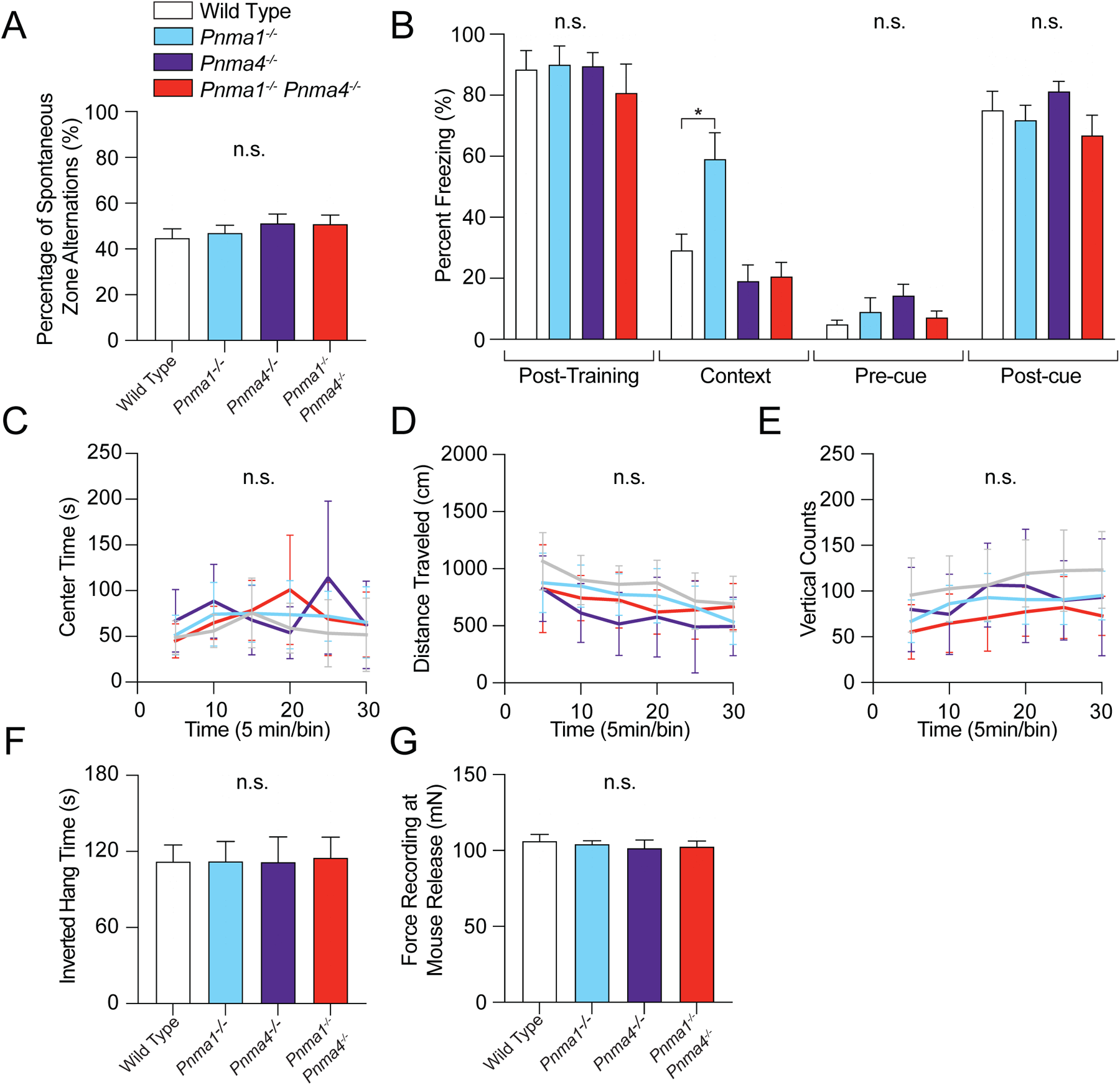
Mice lacking *Pnma1* or *Pnma4* exhibit normal neurobehavioral and muscular traits. Six-month-old male wild type (N=11), *Pnma1^-/-^* (N=11), *Pnma4^-/-^* (N=8), and *Pnma1^-/-^*; *Pnma4^-/-^* double mutant mice (N = 11) were tested in a panel of behavioral assays. **(A)** Short-term memory was assessed using the spontaneous alternation percentage in a Y maze. **(B)** Long-term memory was assessed with the fear conditioning test. **(C-E)** The open field test was used to assess indicators of anxiety (center time) and hyperactivity (**D**, distance travelled and **E**, vertical counts). **(F)** We assessed muscle strength by measuring inverted hang time on a grid and **(G)** grip strength force. All graphs indicate mean +/-SEM. Statistical significance (*p < 0.05) was assessed by multiple t-test analysis with correction for multiple comparisons or one-way ANOVA with correction for multiple comparisons.

**Figure S9:**
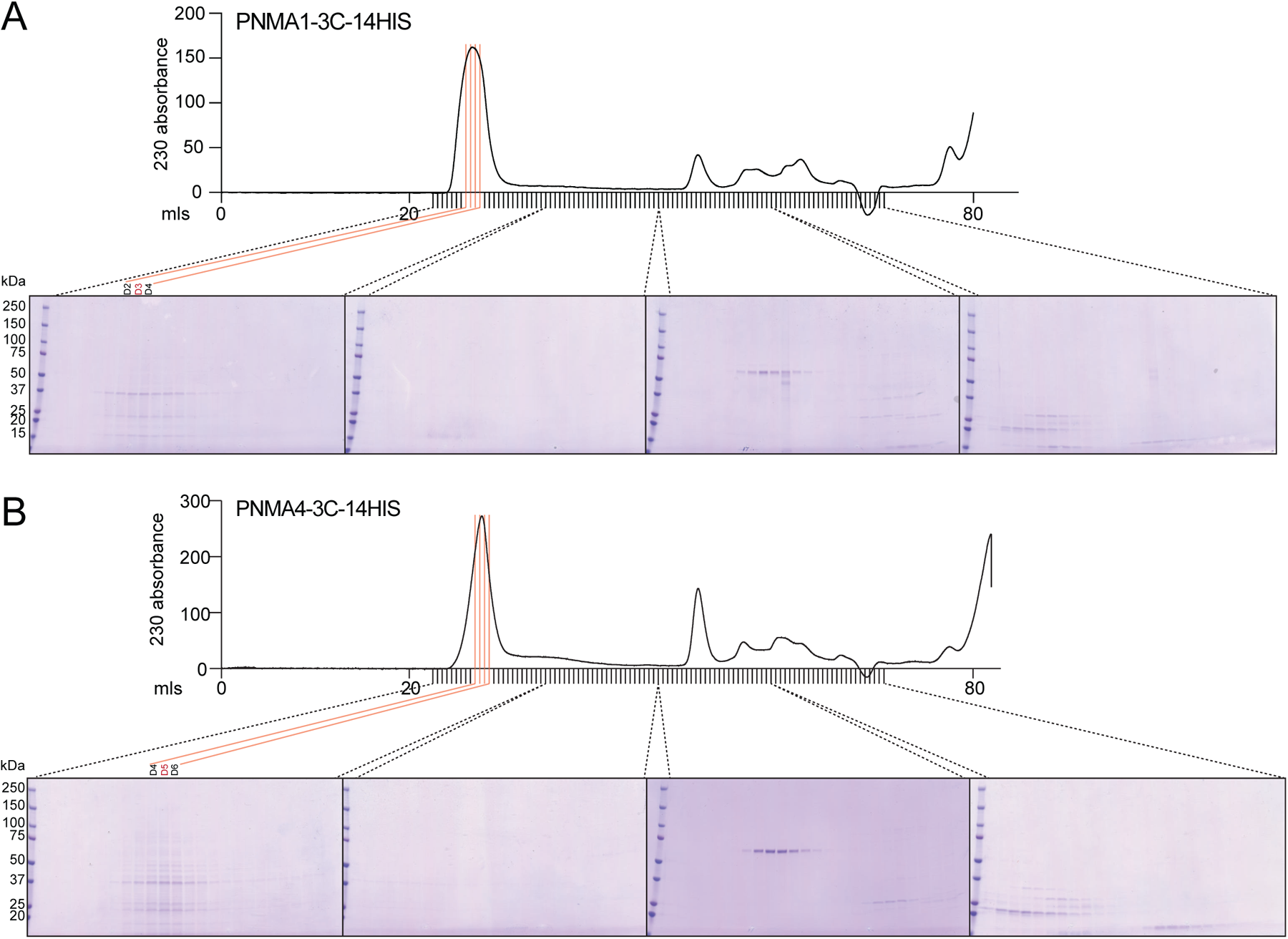
Size exclusion chromatography of recombinant PNMA1 and PNMA4. **(A, B)** His-tagged PNMA1 (A) and PNMA4 (B) were expressed in *E. coli* and affinity purified on nickel resin. Imidazole elutions (15 ml) were concentrated and run over a Superose 6 (HiScale 80 ml) size exclusion column. Shown are UV absorbance traces of the run and Coomassie-stained SDS-PAGE of the fractions. The fractions used for TEM are noted in red.

**Figure S10:**
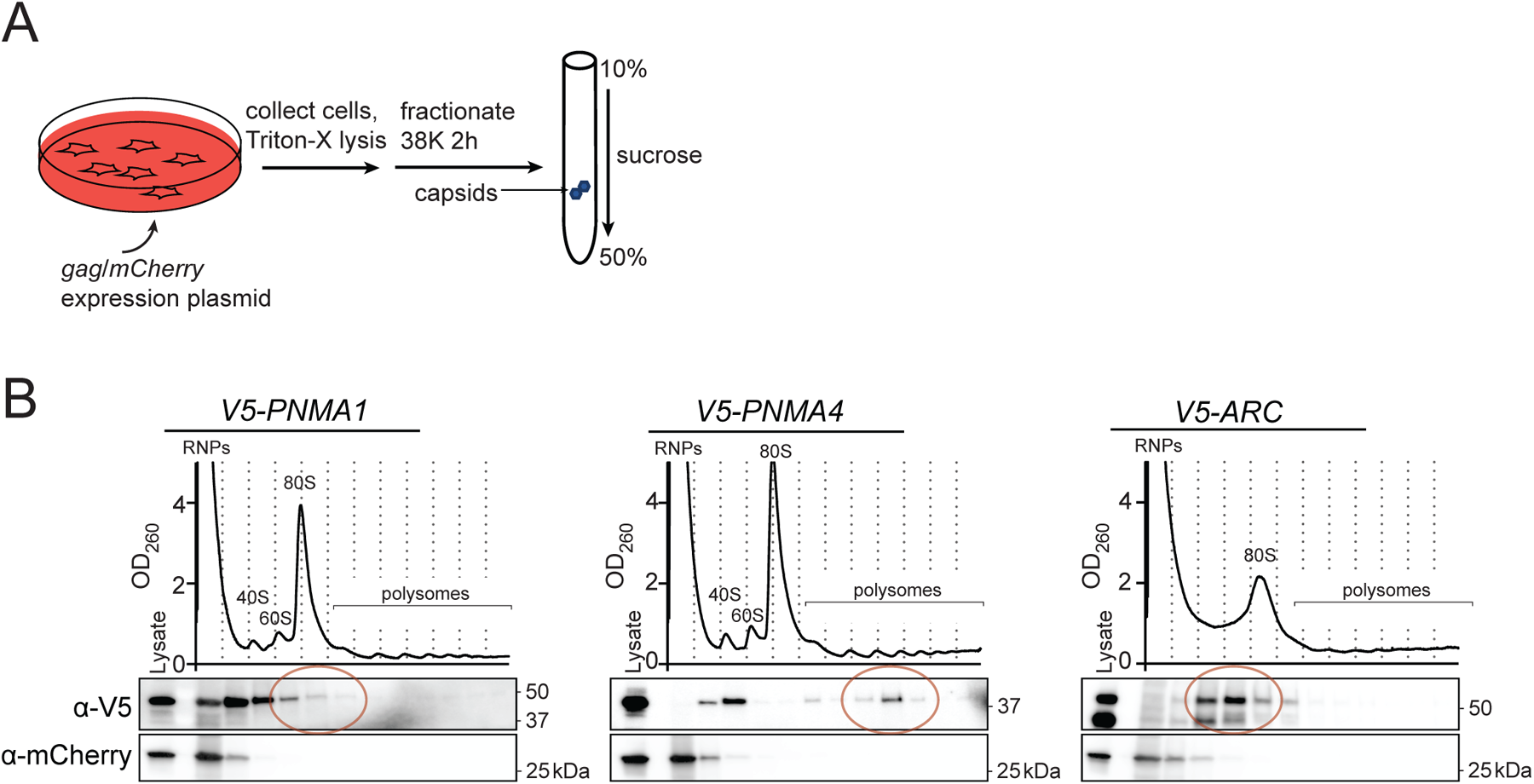
PNMA1 and PNMA4 expressed in human cells forms capsid-sized particles. **(A, B)** Experimental setup (A): *3V5*-tagged *PNMA1*, *PNMA4*, and *ARC* (control exported VLP) expression plasmids were transfected into HEK-293T cells. *mCherry* (control non-capsid) was co-expressed from the transfected plasmid. Cells were collected and lysed in Triton-X. Lysates were fractionated on 10-50% sucrose density gradients with continuous monitoring at 260 nm. PNMA1, PNMA4, ARC (anti-V5), and mCherry protein levels in each fraction were analyzed by immunoblot.

**Figure S11:**
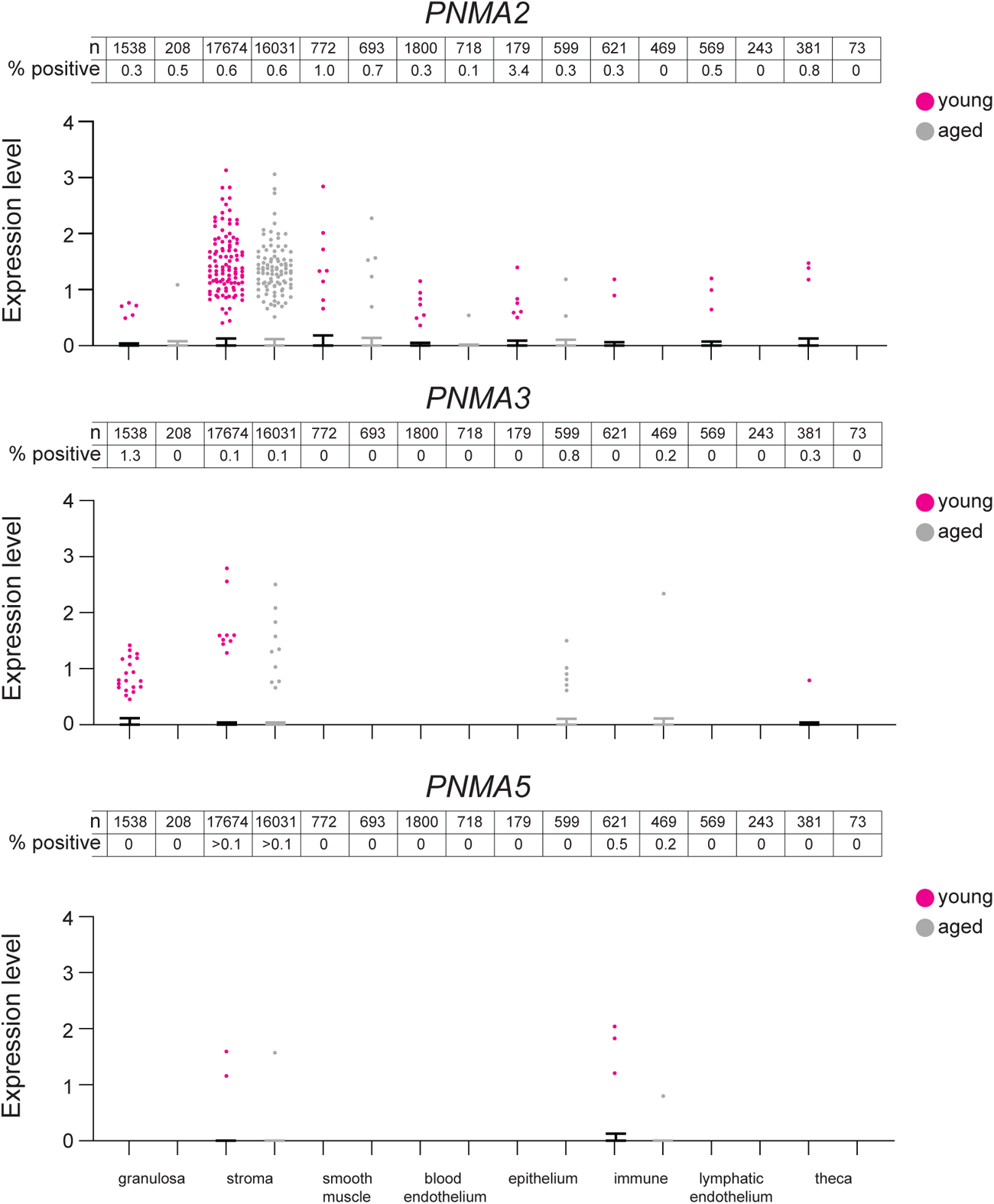
*PNMA2*, *PNMA3*, and *PNMA5* expression analysis in human ovaries. Ovaries from reproductively young (23-29 years, magenta, N = 4) and reproductively old (49-54 years, gray, N = 4) donors were analyzed by single-nuclei RNAseq. Uniquely mapping reads for *PNMA2*, *PNMA3*, and *PNMA5* loci were assigned to ovarian tissue types based on clustering analysis. Statistical significance (*p < 0.05) was determined by Mann-Whitney test.

**Figure S12:**
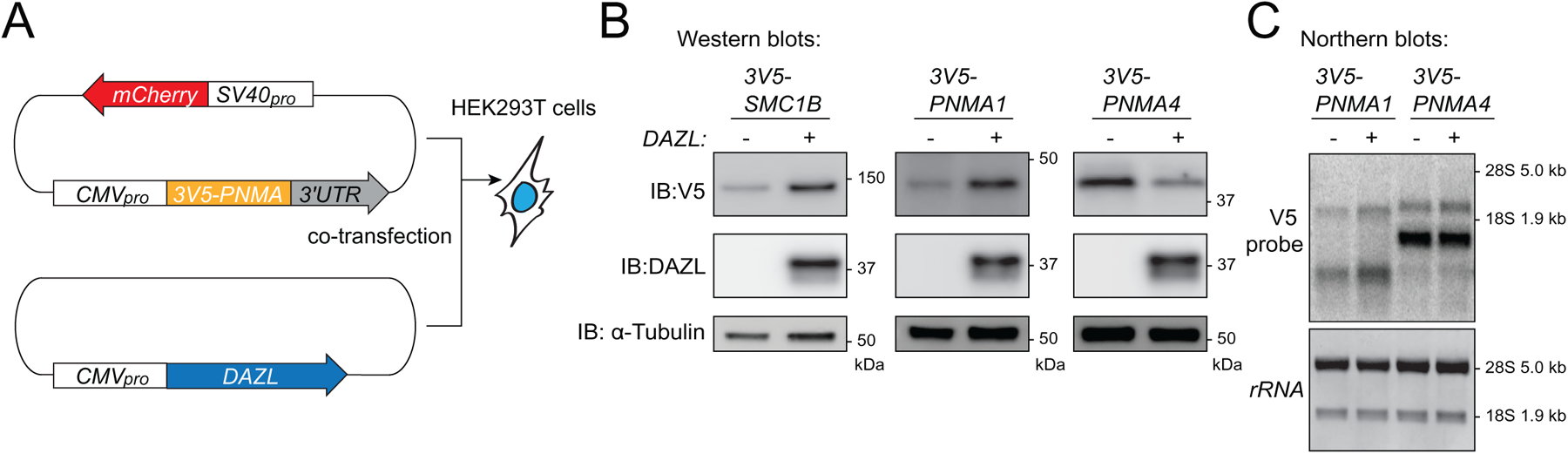
DAZL has the capacity to post-transcriptionally regulate *PNMA1* and *PNMA4*. **(A)** 3V5-tagged *PNMA1*, *PNMA4*, or *SMC1B* (positive control DAZL target) expression plasmids were transfected into HEK-293T cells with and without *DAZL* co-transfection. **(B)** Protein levels of PNMA1, PNMA4, SMC1B, DAZL, and α-tubulin (loading) were analyzed by immunoblot. **(C)** *PNMA1* and *PNMA4* mRNA levels were analyzed by Northern blot.

## METHODS

### Generation of mutant mouse models

*Pnma1^-/-^* and *Pnma4^-/-^* mice were generated at the Columbia University Irving Medical Center (CUIMC) Genetically Modified Mouse Model Shared Resource. Two single guide RNAs (sgRNA) were assembled with the Alt-R™ S.p. Cas9 Nuclease V3 (IDT) in an injection buffer containing 5 mM Tris-HCl [pH 8] and 0.2 mM EDTA. This assembly process aimed to form the CRISPR RNP (ribonucleoproteins) with a final concentration of 0.1 µM. The assembly was conducted at room temperature for 10 minutes. Subsequently, the assembled RNP was microinjected into the pronuclei of fertilized B6CBAF1 eggs. Eggs that survived the injection were then transferred into the oviducts of E0.5 pseudo-pregnant surrogates. Pups resulting from this injection were genotyped at the age of 15 days using PCR.

Along with these knockout lines, the following mouse strains were used in this work: C57BL/6J (wild type Jackson Labs: #000664), B6D2F1/J (Jackson Labs: #100006), and CF-1 (Inotiv/Envigo: 033-US).

### Mouse husbandry

Animal studies were conducted in accordance with the Institutional Animal Care and Use Committee (IACUC) policies of CUIMC and the respective NIH policies. All studies were also conducted in accordance with the provisions of the Animal Welfare Act (NIH/DHHS) and the guidelines of the Association for Assessment and Accreditation of Laboratory Animal Care (AAALAC). All mice were bred at CUIMC, the Jackson Laboratory, or Inotiv/Envigo Animal facilities. Mice were housed at CUIMC. All mouse procedures were conducted at CUIMC except for oocyte dissection, which was conducted at Rutgers University and Yale University. Mice were checked daily, housed in ventilated temperature-controlled cages that were regularly cleaned, containing no more than 5 individuals, and with *ad libitum* access to food and water. Mice were given ample time to equilibrate to housing facilities/experimental rooms as necessary. The animal protocols for Columbia University were: AABO0557 (L.E.B.), AABM7575 (V.A.G.). For Rutgers University: PROTO201702497 (K.S.). For Yale University: 2021-20408 (B.M.). Along with knockout mouse lines created at CUIMC mentioned above, the following mouse strains were used in this work: C57BL/6J (wild type Jackson Labs: #000664), B6D2F1/J (Jackson Labs: #100006), and CF-1 (Inotiv/Envigo: 033-US). The ages of mice are detailed in the respective sections of the manuscript.

### Mouse genotyping

Mouse tail clips were added to 200 µl of 50 mM NaOH in a 1.5 ml tube and placed in a thermomixer at 350 rpm at 55°C overnight. The next day, 25 µl of 1 M Tris-HCL [pH 7.5] was added. Samples were then vortexed and spun at 16,100*g* to pellet debris. Supernatant was transferred to a new tube and frozen at -20°C until genotyping. For PCR we used the following primers:

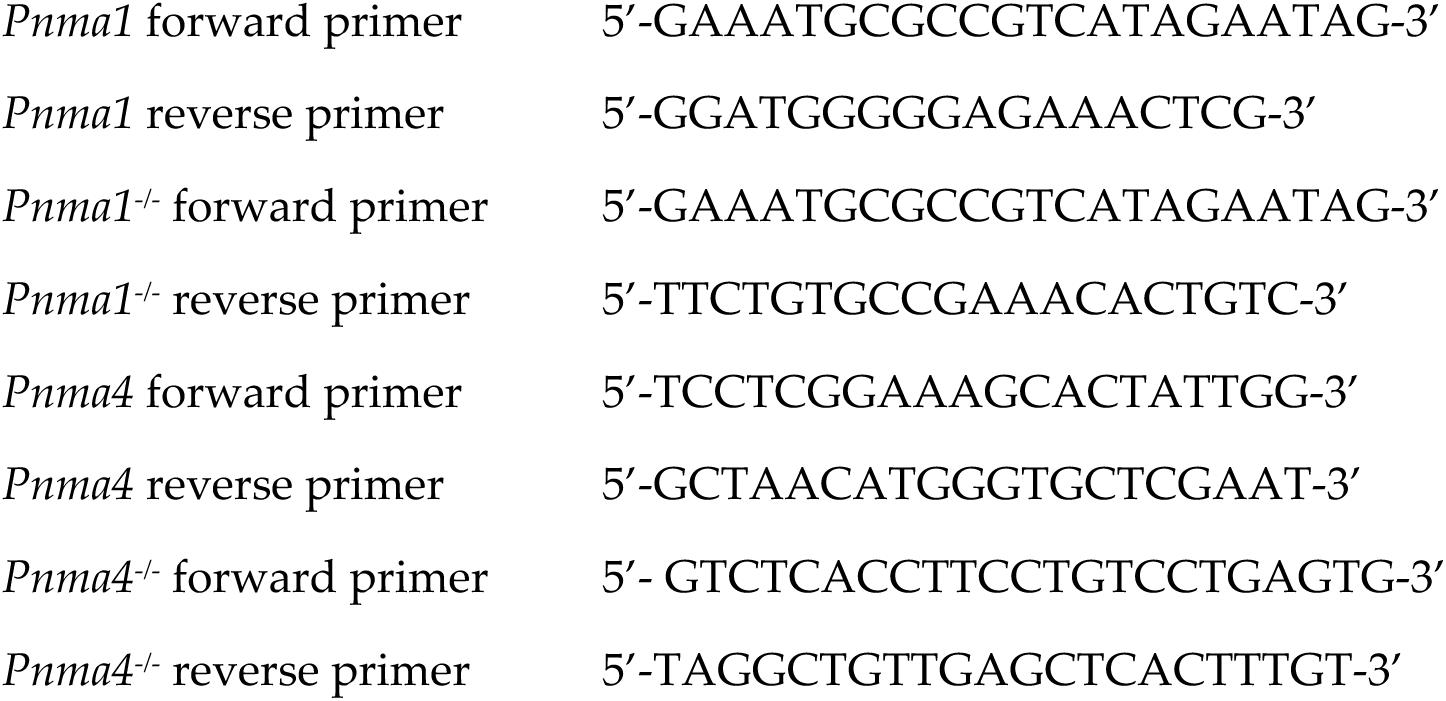

Standard thermocycler amplification conditions were used with Platinum SuperFi II DNA Polymerase and corresponding buffer (Thermo Fisher, REF-12361050), an annealing temperature of 60°C, and 35 cycles. In early rounds of genotyping, following amplification, bands were cut from the gel and extracted using column purification (Macherey-Nagel, REF-740609.250). Samples were submitted for sequencing at Azenta/Genewiz to confirm gene deletions or fragment insertions.

### Fertility assay

To assess mouse fertility, *Pnma1^-/-^, Pnma4^-/-^*, or C57BL/6J wild type control mice were crossed to fertility tester mouse strains for 7 (females) or 14 (males) breeding cycles beginning at ∼2 months of age. One male was paired with one female in a cage at a time. Female *Pnma1^-/-^, Pnma4^-/-^*, or C57BL/6J wild type mice were crossed to B6D2F1/J male tester mice. Mice were paired for a week and then separated. Males were then reintroduced three weeks after being separated (four weeks after the beginning of the prior cross). Pups were counted, removed, and sacrificed before reintroducing mice. Male *Pnma1^-/-^, Pnma4^-/-^*, or C57BL/6J wild type mice were crossed to CF-1 female tester mice. Mice were paired for a week and then males separated from female group ‘A’, with males being paired with another female ‘B’ after a week of separation from female ‘A’. Males were paired with female ‘B’ for a week, separated for a week and then returning to female ‘A’ and repeated. This was done to provide more data for male fertility studies. Pup counts from female and male fertility trials were recorded and plotted with age of parents at the time of the cross. Statistical significance (*p < 0.05) was determined by student’s t-test.

### Body and gonad weight

Mouse body weight was measured prior to dissection using an analytical balance (Mettler Toledo, MS104TS/00). Ovaries and testis were weighed with as much fat and connective tissue removed as possible without damaging the morphological integrity of the organs.

### Immunofluorescence of testis sections

Testes were dissected and placed into 3 ml of 4% PFA (16% stock, Electron Microscopy Sciences, REF-15710) in a 15 ml falcon tube. Testes were incubated in fixative at 4°C with rocking overnight. The following morning, samples were washed 3x (20 minutes) in 1x PBS (10x stock, Fisher Bioreagents, REF BP3994) at 4°C with rocking. Samples were washed ∼10 x in 70% EtOH at 4°C with rocking. Samples were submitted in 70% EtOH to the Columbia University Medical Center Molecular Pathology Core for paraffin embedding and slides containing 5 µm sections were prepared by the Rutgers Research Pathology Services Core Facility. Slides were deparaffinized by 3 x 5-minute washes in xylene, followed by rehydration by 3-minute washes in 100% EtOH, 95% EtOH, 70% EtOH and DI water.

TUNEL assays were performed using the Click-iT Plus TUNEL Assay kit (Invitrogen, C10617) according to manufacturers’ instructions with the following modifications: after the TdT reaction, detection was done using 1/10^th^ of the recommended Click-iT Plus TUNEL reaction cocktail. For immunofluorescence staining of testis sections, antigen retrieval was performed by treating slides for 20 minutes in Sodium Citrate buffer (10 mM Sodium Citrate, 0.05% Tween 20 in Milli-Q water, pH 6.0) at 95-100°C. Slides were blocked for 20 minutes in blocking buffer (0.2% BSA, 0.2% gelatin, 0.05% Tween 20 in PBS), incubated with goat anti-PLZF primary antibody at 1:500 dilution (Human PLZF Antibody, R&D Systems AF2944) at 4°C overnight. Stained slides were blocked by 3 x 10-minute washes in blocking buffer, incubated with donkey anti-goat Alexa Fluor 488 at 1:250 dilution (Invitrogen A-11055) for 60 minutes at 37°C, and blocked by 3 x 10-minute washes in blocking buffer. Slides were mounted using mounting medium containing DAPI (Vectashield). Slides were digitized using EVOS M700 Imaging System (Invitrogen) with a 40× objective and analyzed using QuPath software^84^. Images in figures were captured using a Leica TCS SP8 STED confocal microscope. Images of PLZF-staining were subjected to wavelet-based noise reduction using Leica LAS X software.

### Epididymal sperm counts

To assess epididymal sperm count in mice, the epididymis was separated from the testis. The cauda epididymis and deferent duct were then dissected and washed in 1 ml of 1x PBS (10x stock, Fisher Bioreagents, REF-BP3994) in a plastic petri dish. The cauda epididymis and deferent duct were then placed in 0.5 ml of 10 mM sodium citrate [pH 6.0] in a plastic petri dish and shredded with needles to release sperm. Epididymal tissue was allowed settle onto the dish prior to the removal liquid supernatant to a 1.5 ml tube. Additional sperm was collected from debris by washing with 0.5 ml of 10 mM sodium citrate [pH 6.0] for 1 ml total suspension. Sperm suspension was then diluted 1: 5 with 10 mM sodium citrate [pH 6.0]. Samples were stored at 4°C before counting. 10 µl of diluted sperm suspension was loaded onto a hemocytometer and five regions were counted. This counting process was repeated 3 times with a fresh diluted sperm sample. To calculate the number of sperm in one sample, the average of the counts was multiplied by 10,000 to account for the dilutions.

### Periodic acid-Schiff (PAS) staining

Organs were dissected into 3 ml of Bouins solution (Sigma, REF-HT10132-1L) in a 15 ml falcon tube and incubated at 4°C with rocking overnight. The following morning, samples were washed 3 x 20 in MilliQ water at 4°C with rocking. Samples were then washed ∼10 x in 70% EtOH at 4°C with rocking, or until the EtOH was clear. Samples were submitted in 70% EtOH to the CUIMC Molecular Pathology Core for paraffin embedding, sectioning, and PAS staining.

### Blood collection, sample preparation, and hormone analysis

Mice were anesthetized via isoflurane and sacrificed by cervical dislocation followed by decapitation. 0.5 ml-1 ml of blood was collected into a 1.5 ml Eppendorf tube. Blood was allowed to clot at room temperature for 90 minutes and then the clot was disrupted from adhesion to the tube wall with a wooden applicator stick. Samples were centrifuged at 2000*g* for 15 minutes at room temperature to separate serum from blood. Serum was pipetted into a fresh 1.5 ml Eppendorf tube and frozen at -20°C until shipment for analysis. Samples were shipped overnight on dry ice to the University of Virginia Ligand Core for hormone analysis.

### Ovarian histology

PAS-stained ovary sections were imaged using an EVOS XL core microscope at 10 x magnification. Images were analyzed using FIJI/ImageJ software. Total area was measured via the freehand selections tool and numbers of ovarian features were quantified including antral follicles, secondary follicles, primary follicles, primordial follicles, corpora lutea, post ovulation follicles, follicular cysts, and instances of follicular atresia. Variability of section size was accounted for by calculating the prevalence of each feature per mm^2^.

### Oocyte collection and maturation

Oocytes arrested in meiotic prophase I were collected from ovaries of 6-month-old females as described previously. Briefly, ovaries were placed in minimal essential medium (MEM) containing 2.5 µM milrinone (Sigma-Aldrich #M4659) to prevent meiotic resumption. Oocytes were isolated by piercing the ovaries with needles. To induce meiotic resumption, oocytes were cultured in milrinone-free Chatot, Ziomek, and Bavister (CZB) medium in an atmosphere of 5% CO_2_ in air at 37°C. Oocytes were matured for 16 hours to reach metaphase II, fixed in phosphate buffer saline (PBS) containing paraformaldehyde (PFA) 2% for 20 minutes at room temperature, and permeabilized in PBS containing 0.1% (vol/vol) Triton X-100 and 0.3% (wt/vol) BSA for 20 minutes, and blocked for 10 minutes in blocking buffer (0.3% BSA containing 0.01% Tween in PBS). Immunostaining was performed by incubating cells in Alexa-fluor 488 conjugated primary ⍺-tubulin antibody (1:100; Life Technologies #322588) for 1 hour in a dark, humidified chamber at room temperature followed by 3 consecutive 10-minute incubations in a blocking buffer. After washing, the cells were mounted in 10 µL VectaShield (Vector Laboratories, #H-1000) with 4′, 6-Diamidino-2-Phenylindole, Dihydrochloride (DAPI; Life Technologies #D1306; 1:170).

Oocytes were imaged using a Leica SP8 confocal microscope equipped with a 40X, 1.30 N.A. oil immersion objective. For each image, optical z-slices were obtained using a 1.0 µm step with a zoom setting of 2.5. Numbers of metaphase II eggs were recorded.

### Mouse oocyte isolation, culturing, microinjection and high-resolution confocal microscopy

All animal work for visualizing meiosis in live oocytes was conducted in accordance with a protocol approved by the Yale University Institutional Animal Care and Use Committee (IACUC number 2021-20408). Wild-type C57BL/6 female mice were purchased from The Jackson Laboratory. Oocytes were isolated from ovaries of wild-type and *Pnma* knockout mice and cultured in M2 medium containing dbcAMP (Sigma, D0627-100MG) to maintain them in prophase arrest. mRNA transcription constructs for synthesis of MAP4 (to mark microtubules) and H2B (to mark chromosomes) were generated as described previously^85^. Oocytes were microinjected with 6 to 8 pl of *in vitro* transcribed mRNAs as described in detail recently^86^.

For live imaging of meiotic spindle assembly and chromosome segregation, oocytes were released from prophase arrest by transferring them into M2 medium that did not contain dbcAMP. Confocal timelapse images were acquired using Zeiss LSM 800 and Zeiss LSM 900 microscopes equipped with 40× C-Apochromat 1.2 numerical aperture (NA) water-immersion objectives and environmental chambers maintained at 37°C. Image acquisition using ZEN blue software (Zeiss) was performed at a temporal resolution of 6 min and with a Z-stack thickness of 40.5 µm at 1.5 µm confocal sections.

### Plasmid creation

PCDNA4TO (Thermo Fisher, REF-V102020) was provided by Dr. Yossi Sabo. *DAZL* (human) gene fragments were synthesized by Twist Biosciences and cloned into the otherwise unmodified PCDNA4TO. For *PNMA1, PNMA4* (both human and mouse), *SMC1B* and *ARC* plasmids we modified PCDNA4TO by inserting an *mCherry* reporter driven by the secondary SV40 promoter to act as a transfection efficiency reporter and a monomeric protein control. 3V5 tagged *Pnma1* and *Pnma4* (mouse) and *PNMA1, PNMA4*, *SMC1B* and *ARC* (human) gene fragments were synthesized by Twist Biosciences and cloned into the modified PCDNA4TO backbone in the Berchowitz lab using standard restriction cloning procedures. Plasmid sequences were analyzed using Genewiz/Azenta.

### HEK293T Cell Culture

HEK293T cells were grown on 6-well flat bottom with lid culture plates (tissue culture treated, non-pyrogenic, polystyrene, Corning, REF-3516). Cells were maintained in a grown medium consisting of Dulbecco’s Modified Eagle Medium (DMEM) +4.5 g/L D-Glucose, - sodium Pyruvate (Gibco, REF-11960-044), supplemented with 10% Heat Inactivated Fetal Bovine Serum (HI-FBS) (Sigma, REF-F4135), 1% L Glutamine 200 mM (Gibco, REF-25030-081), and 1% penicillin-streptomycin (Sigma, REF-P4458). Cells were incubated at 37°C, with 5% CO_2_ in a humidified environment (HERAcell 150i, ThermoFisher Scientific). Cells were passaged when they reached ∼80% confluency after ∼2 days and reseeded at 5 x 10^5^ cells per well of a 6-well plate. During passaging, cells were dissociated using 0.25% Trypsin-EDTA, phenol red (ThermoFisher Scientific, REF-25200056) for 2 minutes at 37°C and inactivated via dilution with un-supplemented DMEM. Cells were centrifuged at 300 RCF for 5 minutes and resuspended in supplemented medium and then reseeded in the necessary dishes.

### HEK293T Transfection

On experimental day one, cells were passaged into a 24-well plate (2.5x10^5^ cells per well) with 450 µl of supplemented DMEM and incubated for 24 hours as described above. On experimental day 2, old media was aspirated and replaced with premixed 250 µl supplemented DMEM and 200 µl OPTI-MEM (Gibco, REF-31985-070) and then placed back in the incubator. To prepare transfection mix tubes: tube 1 (general tube) contained 24 µl per well of OPTI-MEM, and 1ul per well of Lipofectamine 3000 (Thermo Fisher, REF-L3000008), tube 2 (well specific tube, 2A, 2B, etc.) contained 1 µl of P3000 enhancer (Thermo Fisher, REF-L3000008), 1 µg/µl of plasmid DNA, and 23 µl of OPTI-MEM. We added 25 µl of tube 1 to each tube 2 and incubated them in the tissue culture hood for 10-15 minutes at room temperature. The 50 µl of transfection mix was added to the wells using gradual dropwise pipetting, gently rocked to distribute transfection mixture, and cells were placed back in the incubator for 24 hours. On experimental day 3, cells were checked for transfection efficiency using fluorescent reporters (Evos M5000). If transfection was successful, cells were dissociated as described above and prepared for western/northern blotting or density gradient ultracentrifugation analysis.

### Cloning, expression, and purification of recombinant PNMA1 and PNMA4

Full-length mouse *Pnma1* and *Pnma4* (DNA fragments synthesized by Twist Biosciences) were cloned into the first cassette of a pET-Duet plasmid using inverse PCR. The pET-Duet plasmid was previously modified to encode a C-terminal 3C protease cut site) followed by a His tag. For protein expression, the plasmids carrying C-terminally tagged *Pnma1* and *Pnma4* were transformed into the LOBSTR *E. coli* expression strain containing a RIL plasmid (Kerafast)^87^. To obtain purified PNMA1 and PNMA4, 1L of LOBSTR cells containing the *Pnma1* or *Pnma4* plasmids were grown in LB (with 100 µg/mL of ampicillin and 35 µg/mL of chloramphenicol) to an OD_600_ of 0.7 at 37°C, shifted to 18°C for 30 min, and induced with 200 µM IPTG for 18 hrs. The cells were collected by centrifugation and resuspended in 400 mL of cold lysis buffer (50 mM Tris/HCl pH 8.0, 500 mM NaCl, 40 mM imidazole, and 5 mM β-mercaptoethanol). The cells were lysed on ice by sonication (Misonix). Cell debris was removed by centrifugation and the supernatant was incubated with 1 mL of equilibrated Ni-Sepharose 6 Fast Flow resin (GE Life Sciences) and gently mixed in batch for 1 hr at 4° C. The Ni-resin was collected by centrifugation and placed into a 10 mL polypropylene column (Thermo Scientific) and washed with 100 column volumes of wash buffer (10 mM Tris/HCl pH 8.0, 150 mM NaCl, 40 mM imidazole, and 5 mM β-mercaptoethanol). Pnma1 and Pnma4 were eluted with 10 mL of elution buffer (10 mM Tris/HCl pH 8.0, 150 mM NaCl, 300 mM imidazole, and 5 mM β-mercaptoethanol) and analyzed by SDS-PAGE. Samples were then concentrated over a 30 kD molecular weight cutoff (MWCO) filter to ∼500 µl, and run over a Superose 6 Increase HiScale (80 ml volume, Cytiva) size exclusion column. Fractions were analyzed by SDS-PAGE (**Figure S9**) Finally, the void fractions corresponding to large forms of PNMA1 and PNMA4 were collected for further analysis.

### Negative stain transmission electron microscopy

Aliquots of PNMA1 and PNMA4 were thawed on ice and centrifuged for 5 minutes at 4° C at 16,000*g* to pellet any insoluble protein. Samples for negative stain TEM were prepared as described^88^. Formvar/Carbon 300 Mesh, Cu grids (Electron Microscopy Sciences, FCF300-Cu-50) were glow discharged using 15 mA for 45 s, with a 10 s hold (easiGlow, PELCO). 3 µL (PNMA4) or 5 µL (PNMA1) of sample was spotted on glow-discharged grids at room temperature. After 1 minute, sample was blotted away, washed twice with water, and stained with 2% uranyl acetate (made from SPI supplies, CAS#6159-44-0). Samples were dried for at least ten minutes prior to imaging. Images were acquired using DigitalMicrograph from Gatan at either 15000 or 23000 nominal magnification and converted to MRC files in DigitalMicrograph.

Micrographs were imported into cryoSPARC^89^ (v4.4.1+240110) (Output constant CTF: true). Capsid-like particles were manually picked and classified using the 2D classification job (Re-center 2D-classes: false, Do CTF correction: false). For PNMA4, the partial staining observed prevented use of the template picker. However, for PNMA1, the complete negative staining allowed us to use the template picker followed by several rounds of 2D classification to identify capsid-like structures.

### Velocity gradient ultracentrifugation

HEK293T cells were transfected with 3*V5*-tagged *PNMA1*, *PNMA4*, or *ARC* expression plasmids as described above. Approximately ∼4 x 10^6^ cells were lysed in polysome lysis buffer (1% Triton-X, 20 mM Tris-HCl [pH 7.5], 10 mM magnesium chloride, 50 mM potassium chloride, 10 µg/mL cycloheximide, 1 mM PMSF, 1x Halt protease and phosphatase inhibitor cocktail (Thermo Fisher Scientific, 78442)). Lysate was cleared by centrifugation at 4°C at 20,000*g* for 10 minutes. Lysate was loaded on a 10% to 50% sucrose gradient in polysome lysis buffer which was centrifuged at 4° C for 3 hrs at 30,000 rpm in a Beckman SW41Ti rotor. Fractions were collected using a BioComp gradient station and a BioComp Triax flow cell monitoring continuous absorbance at 260 nm. For Western blot analysis, each fraction received 100% TCA to a final concentration of 5% TCA. Precipitate was pelleted by centrifugation and washed once in acetone. We added 99 µl TE, 1 µl 1M DTT, and 50 µl Loading Buffer (9% SDS, 0.75 mM Bromophenol blue, 187.5 mM Tris-HCl [pH 6.8], 30% glycerol, and 810 mM beta-mercaptoethanol); samples were heated at 100°C for 5 min and centrifuged 5 min at 20,000 g prior to analysis by SDS-PAGE.

### Western Blotting

To prepare testes lysate (for protein and RNA extraction), one snap-frozen testis was ground by mortar and pestle into a fine powder under liquid nitrogen. 0.75 ml lysis buffer (150 mM NaCl, 50 mM Tris [pH 7.5], 0.25% NP-40, 1 mM PMSF, 1x Halt Protease Inhibitor Cocktail (Pierce)) was added. The lysate was clarified by centrifugation at 20,000 g for 20 minutes, collected to a new tube, and re-clarified by centrifugation at 20,000 g for 20 minutes. 100 µl lysate was combined with 50 µl 3x SDS loading buffer (9% SDS, 0.75 mM Bromophenol blue, 187.5 mM Tris-HCl [pH 6.8], 30% glycerol, and 810 mM β-mercaptoethanol) and samples were boiled at 100°C for 5 minutes, and then centrifuged at 20,000 x g for 5 minutes. To prepare HEK293T lysate, 5x10^5^ cells were resuspended in 100 µl lysis buffer (85 µl RIPA buffer (Sigma, REF-R0278), 10µl TURBO DNase buffer (ThermoFisher Scientific, REF-AM2238), 2 µl TURBO DNase (ThermoFisher, REF-AM2238), 1 µl RNaseA at 5 µg/µl (Lucigen, REF-MRNA092), 1 µl HALT protease inhibitor (ThermoFisher Scientific, REF-1861282), 1 µl PMSF (ThermoFisher, REF-36978)) and incubated at room temperature for 1 minute. 50 µl 3x SDS loading buffer was added, samples were boiled at 100°C for 5 minutes, and centrifuged at 20,000*g* for 5 minutes.

Pre-cast polyacrylamide gels (4%-15% Mini-Protean TGX Precast Gels, Bio-Rad, 10 well REF-4561083, 15 well REF-4561086) were loaded and run on a mini gel system (Mini-PROTEAN Tetra Vertical Electrophoresis Cell, Bio-Rad REF-1658004) at 200V for 35-45 minutes. Gels were run with precision plus protein dual color standards (Bio-Rad, REF-1610374) and with SDS Running Buffer (190 mM glycine, 25 mM Trizma base, 3.5 mM, 1% SDS). Transfers to nitrocellulose (Bio-Rad, REF-1620145) were performed using semi-dry transfer on a trans turbo blot transfer apparatus (Bio-Rad, REF-1704150), with turbo transfer filter paper for nitrocellulose (Bio-Rad 1620215) and stained in Ponceau S to visualize total protein load. Membranes were blocked with 3% milk powder in TBST for 1 hour. Primary antibody solutions were prepared in 50 ml of 1x TBST, 2% Sodium azide, 1% milk 1% BSA plus antibody at necessary concentration. Membranes were incubated with primary antibodies at 4°C with rocking overnight. The next day, primary antibody solution was removed, and the membrane given 2x 10-minute 1x TBST washes. Secondary antibody solutions were prepared in 50 ml of 1x TBST, 1% milk 1% BSA plus antibody at necessary concentration. After ∼2 hours (or longer if necessary) secondary antibody solutions were removed from membranes, which were washed 2x (10-minute 1x TBST) before visualization.

Primary antibodies were used at the following concentrations: α-PNMA1 1: 2,000 (Proteintech 13631-1-AP), α-PNMA1/MOAP1 1: 2,000 (Proteintech 15325-1-AP), α-α-tubulin 1: 1,000 (Invitrogen), α-V5 1: 1,000 (Invitrogen), α-DAZL 1: 5,000 (Abcam), α-mCherry 1: 2,000 (Abcam), α-GAPDH 1:5,000 (Thermo Fisher), and α-MLV p30 1: 1,000 (Abcam). α-mouse HRP-conjugated, α-rabbit HRP-conjugated (Cytiva), or α-rat FITC-conjugated (for α-α-tubulin, Thermo Fisher) secondary antibodies were used.

### Quantitative PCR

Total RNA was prepared from 400 µl testis lysate prepared as in previous section. SDS was added to 0.5%, 400 µL acid phenol: chloroform 5: 1 (Ambion) was added, and samples were incubated at 1,400 rpm for 30 min at 65°C in a Thermomixer (Eppendorf). Samples were centrifuged for 5 min at 13,000*g*, extracted to 1 mL 100% ethanol and 40 µL sodium acetate [pH 5.5], and precipitation was conducted at 20°C overnight. RT and qPCR was conducted using the Primetime one-step RT-qPCR master mix (IDT) with 80 ng RNA template, 500 nM forward/reverse primer, and 250 nM probe. Reactions were conducted in triplicate (with no RT and no RNA controls) in a CFX96 thermocycler (BioRad) using the following parameters: 50°C 15:00 (1x), 95°C 3:00 (1x), 95°C 0:05, 60°C 0:30 (40x).

> ***Pnma1:*** F primer 5’-GAATGGCAGGTGTCGGATATAG-3’ R
>
> primer 5’-GCGTTGTTGGTCTTGAGGAT-3’
>
> probe 5’- /56-FAM/CGGCGGTTG/ZEN/ATGGAGAGTTTGAGA/3IABkFQ/-3’
>
> ***Pnma4:*** F primer 5’- GGACCAGGTGAGGAAGAATTT-3’ R
>
> primer 5’- TGGACCTCTAAGGCTCTCTATC-3’
>
> probe 5’-/56-FAM/ACATGGCAG/ZEN/GTGTCAGATGTAGAGA/3IABkFQ/-3’
>
> ***PPIA:*** F primer 5’- GGCTATAAGGGTTCCTCCTTTC-3’ R
>
> primer 5’- TTTCTCTCCGTAGATGGACCT-3’
>
> probe 5’- /56-FAM/ATGTGCCAG/ZEN/GGTGGTGACTTTACA/3IABkFQ/3’

### Extracellular protein analysis

HEK293T cells were transfected with 3*V5*-*PNMA1*, 3*V5*-*PNMA4*, or 3*V5*-*ARC* expression plasmids as described above. 40 ml extracellular medium was collected and filtered (0.45 µm) to remove cells and large debris. Cells were collected and lysed in 85 µl RIPA buffer (cell lysate control, analyzed as above). Growth medium was concentrated over a 100 kD MWCO filter to ∼500 µL which was diluted to 8 ml in 1x PBS and layered on top of a sucrose cushion of 2 ml 25% sucrose in an SW41 tube (Seton). The tubes were spun for 3 hr at 38,000 rpm at 4° C in an SW41Ti rotor. After centrifugation, the following fractions were taken: lipids, medium (from middle of 8 ml), meniscus of 25% sucrose, 25% sucrose, and pellet. A sample of each fraction was taken for analysis by SDS-PAGE/immunoblot.

### Analysis of testicular capsid-like PNMA4 protein

***Lysate preparation:*** 10 snap-frozen testes from wild type (C57BL/6J) mice were ground by mortar and pestle into a fine powder under liquid nitrogen. 1 ml lysis buffer (150 mM NaCl, 50 mM Tris [pH 7.5], 0.25% NP-40, 1 mM PMSF, 1x Halt Protease Inhibitor Cocktail (Pierce)) was added. The lysate was clarified by centrifugation at 20,000 g for 20 minutes, collected to a new tube, and re-clarified by centrifugation at 20,000 g for 20 minutes and diluted to 8 ml in lysis buffer.

***Double sucrose gradient:*** 8 ml testis lysate was layered on top of a double sucrose cushion of 2 ml 25% (upper) and 2 ml 70% (lower) in an SW41 tube (Seton). The tubes were spun for 3 hr at 38,000 rpm 4° C in an SW41Ti rotor. After centrifugation, the following fractions were taken: lipids, lysate (from middle of 8 ml), meniscus of 25% sucrose, 25% sucrose, meniscus of 70% sucrose, 70% sucrose, and pellet. A sample of each fraction was taken for analysis by SDS-PAGE/immunoblot.

***Iodixanol step gradient:*** An iodixanol (Optiprep density gradient medium, supplied at 60% iodixanol) step gradient was prepared in an SW41 tube with the following steps: (top) 2 ml 15% (in 1 M NaCl), 3 ml 30%, 3 ml 40%, 3 ml 50% (bottom). The 70% meniscus fraction (∼500 µL) was layered on top and the tube was spun at 41,000 rpm for 48 hours at 4° C in an SW41Ti rotor. Twelve 1 ml fractions were taken using a pipette. ∼40 µL samples (20 µL 3x SDS loading buffer was added) from each fraction were taken for analysis by SDS-PAGE/immunoblot (30 µL load).

### Northern blot analysis

1 x 10^7^ HEK293T cells were harvested by centrifugation. RNA was extracted using the RNeasy kit (Qiagen, REF-74104) using the manufacturer’s protocol. We added 22 mL denaturing mix (15 mL formamide, 5.5 mL formaldehyde, and 1.5 mL 10x MOPS) to 8 mg total RNA in 8 mL and heated at 55°C for 15 min 20 mL of sample (approximately 5 mg) was resolved on a denaturing agarose gel (1.9% agarose, 3.7% formaldehyde, 1x MOPS buffer) for 2.5 h at 80 V. The gel was blotted to a Hybond membrane (GE Healthcare) by capillary transfer in 10x SSC (1.5 M NaCl, 0.15 M trisodium citrate dihydrate, [pH 7]). The membrane was incubated in a hybridization buffer (0.25 M Na-phosphate [pH 7.2], 0.25 M NaCl, 1 mM EDTA, 7% SDS, and 5% dextran sulfate) at 65°C probed with 32P-labeled V5 DNA probes prepared via Amersham Megaprime DNA labeling kit (GE Healthcare) and Illustra ProbeQuant columns (GE Healthcare), transferred to a phosphor screen, and imaged on a Typhoon imager (GE Healthcare).

### *PNMA1-5* expression analysis in human and mouse ovary

To assess the expression of *PNMA1-5* in the human ovary, we analyzed single-nuclei RNA-seq datasets obtained from eight individuals. These datasets included samples from four younger women aged 23, 27, 28, and 29, and four older women aged 49, 51, 52, and 54. To assess the expression of *PNMA1-5* in the mouse ovary, we performed single cell RNA sequencing of mouse ovaries from reproductive young (Y, 4.5-month, n=5), peri-estropause (M, 10.5-month, n=7), and post-estropause (O, 15.5-month, n=5) stage. The gene expression data and metadata for these samples were downloaded from the GEO database (accession number: GSE202601). Additionally, human oocyte expression data were sourced from two published datasets available under GEO accession numbers GSE107746 and GSE155179. All gene expression data were normalized using the ’NormalizeData’ function from Seurat (v4.4.0)^90^. Previously published data from single cell RNA-seq of adult murine ovarian cells was also used (C57BL/6J mice; n=8). To identify cell-types that express *Pnma1* and *Pnma1,* cells positive for expression of these genes were displayed over a dimplot of previously identified clusters of distinct ovarian cell-types using the featureplot command from the Seurat package. To compare expression of these genes across clusters, the average expression levels from each sample was obtained and used for statistical comparison in the GraphPad Prism software. Strip plots are presented with individual points shown and means ± SEM indicated. For comparisons of means between clusters, one-way ANOVA was used followed by Tukey’s post-hoc test. Significant differences were defined at P < 0.05.

### Analysis of traits associated with human *PNMA1* and *PNMA4* variation

We gathered information on the traits associated with the *PNMA1* and *PNMA4* genes using the Open Targets Genetics database containing GWAS summary statistics^44^. This database predicts the causal genes of traits using the variant-to-gene and locus-to-gene (L2G) pipeline^43^. In the database, we queried the *PNMA1* and *PNMA4* genes to collect the causally-associated traits from the GWAS studies. We then filtered traits by *P*-value < 5x10^-8^ and merged them into a table (**Table S1**) labeled with the queried genes.

### Statistical analysis

All statistical analyses were conducted using Prism (V10, GraphPad). Unless indicated otherwise, statistical significance (*p < 0.05) was determined by student’s t-test (with Welch’s correction for single nuclei RNAseq data), multiple t-tests with correction for multiple comparisons, one-way ANOVA with correction for multiple comparisons, or Mann-Whitney test as indicated. For all figures, the number of individuals is denoted by capital N and number of measurements is denoted by lowercase n.

### Neurobehavioral tests

All mouse behavioral tests were conducted within the CUIMC Neurobehavior Core facility. ∼6-month-old male *Pnma1^-/-^, Pnma4^-/-^*, *Pnma1^-/-^: Pnma4^-/-^* double knockouts or C57BL/6J wild type mice were used for all tests. Mice were given a minimum of 30 minutes to equilibrate to testing rooms before experiments were conducted. To ensure minimal stress for the animals, the tests progressed from least to most stressful (open field, grip strength, inverted hang, Y maze, fear conditioning).

### Open field test

Individual mice were placed into the center of a clear Plexiglass box (27.31 x 27.31 x 20.32 cm), (Med Associates, ENV-510) in a dim room (∼5 lux). Mice were allowed to explore freely for 30 minutes. Mouse behavior, including exploration of different areas of the box, horizontal and vertical locomotion speed and distance, and time spent in the center vs edge were tracked. Tracking was conducted via infrared beams that were embedded along the X, Y, and Z axes of the box and processed by Activity Monitor Version 7 tracking software (Med Associates Inc.). At the end of the test, mice were returned to their home cages and the box was thoroughly cleaned between trials using 70% EtOH.

### Grip strength test

Forelimb grip strength was tested in mice by utilizing a grip strength test meter (BIOSEB). A small metal grid acted as a grasping surface and was connected to the strength meter. The metal grid was positioned so mice would grasp the bottom bar of the grid, with this being parallel to the horizontal plane of the strength meter. The mouse was held by the tail and moved towards the grid until it reached out and grasped the bottom bar of the grid with its forepaws. The mouse’s tail was gradually pulled backwards until the mouse’s torso was horizontal. When the mouse lost its grip, that force recording was taken as the grip strength reading at peak tension. This test was repeated five times for each mouse. The mean of these five trials was recorded.

### Inverted Hang Test

Mice were placed on a grid, which was inverted and placed over a (30 x 30 x 30 cm) plastic box with padding at the bottom. Time was started upon inversion of the grid. Latency to fall was measured for each mouse or after 3 minutes when the trial was ended. At the end of the test, mice were returned to their home cages and the grid was thoroughly cleaned between trials using 70% EtOH.

### Y Maze Spontaneous Alternation

Mice were placed in a plastic Y-maze comprising 3 arms of equal size (30 x 5 x 30 cm) extending from a central area. The test was conducted in a quiet, dim (∼5 lux) room. The rate of spontaneous alternation between arms of the maze was tracked using Ethovision XT software (Noldus) over an 8-minute trial. At the end of the test, mice were returned to their home cages and the maze was thoroughly cleaned between trials using 70% EtOH.

### Fear Conditioning

Fear conditioning studies were performed as previously described^91^. Training and conditioning trials were performed in two identical shock chambers (30 x 24 x 21 cm), which were inside sound-mitigating chambers to limit outside stimuli (Med Associates). Chambers consist of a clear polycarbonate front wall, a white opaque back wall, two steel side walls and a grid floor with a waste collection pan underneath. Chambers were calibrated to deliver shocks of 0.5 milliamps and 80 decibels. A camera on the door of the attenuating chamber was used to record the experimental session. Sessions were analyzed, and % of time freezing was calculated using VideoFreeze software (Med Associates). For the training session, each chamber was illuminated with white house light. An olfactory cue was presented using a of drop vanilla extract on the back wall of the shock chamber. On experimental day 1, mice were placed into the chamber and allowed to explore freely for 2 minutes. After 2 minutes a pure tone lasting 30 seconds was played (5kHZ, 80dB) to serve as the controlled stimulus (CS). In the last 2 seconds a foot shock (0.5 mA) was administered through the floor grid, which served as the unconditioned stimulus (US). Over the course of the day 1 trial, each mouse received three CS-US pairings with 90 second intervals, with the mouse being left in the chamber after the final CS-US pairing for an additional 2 minutes. Percentage of time staying frozen still (freezing behavior) was recorded during this 2-minute interval. The mouse was then returned to their home cage. On experimental day 2 contextual conditioning was tested in the same shock chamber. The same chamber illumination and olfactory cue was present but with no tone or foot shock. Mice were placed in the chamber for 5 minutes with no CS or US delivered. Freezing behavior was scored during this 5-minute period and the mice was returned to its home cage.

On experimental day three cued conditioning was tested. Contextual cues of the cage were changed by covering the grid floor with smooth white plastic and inserting a piece of curved white plastic to cover the side walls. Near infrared light was used in place of white light, and the olfactory cue was changed from vanilla extract to lemon extract. Mice were placed in the chamber and allowed to freely explore for 3 minutes, followed by 3 minutes of the CS tone (5 kHz, 80 dB). During both 3-minute periods, the percentage of time spent freezing was recorded and scored, with the mouse being returned to its home cage. The chamber was thoroughly cleaned between trials using 70% EtOH, with fresh olfactory cue being applied for each new trial.

## ACKNOWLEDGEMENTS

We thank Vicky Brandt, Stephen Goff, and Sang-On Park for guidance and critical reading of the manuscript. We thank Ian Adams and Gerard Karsenty for technical assistance and helpful discussions. We thank the histology cores at CUIMC and at Rutgers, mouse behavior core at CUIMC, the ligand core at University of Virginia, Brent Mayfield, and Jerrin T. George for technical assistance. We thank Chyuan-Sheng Li at the Genetically Modified Mouse Model Shared Resource at CUIMC for generation of knockout mice. We thank Yossi Sabo for plasmids, Chi-Min Ho for preliminary negative stain TEM, Hachung Chung for sharing use of her microscope, and Sam Sternberg for sharing use of his FPLC. Research in the Berchowitz lab is supported by the Schaefer Research Scholars Program, the Irma T. Hirschl Family Trust, and NIH (R35 GM124633 to L.E.B.). Research in the Wiedenheft lab is supported by the NIH (R35 GM134867 to B.W.), the M.J. Murdock Charitable Trust, and the Montana State University Agricultural Experimental Station (USDA NIFA). Research in the Suh lab is supported by the NIH (R01 AG069750 to Y.S.) and a grant GCRLE-1320 from the Global Consortium for Reproductive Longevity and Equality at the Buck Institute, made possible by the Bia-Echo Foundation. Research in the Schindler lab is supported by NIH (R35 GM136340 to K.S.). Research in the Mogessie lab is supported by the Wellcome Trust (213470/A/18/Z) and the NIH (R35 GM146725 to B.M.). Research in the Jain lab is supported by the NIH (R35 GM147130 to D.J.) and Human Genetics Institute of New Jersey laboratory startup funds. Research in the Gennarino lab is supported by the NIH (NINDS R01 NS109858 to V.A.G.), the Neurodegeneration Challenge Network/Chan Zuckerberg Initiative, the Paul A. Marks Scholar Program, the TIGER Award at CUIMC, and the Columbia Stem Cell Initiative. Research in the Stout lab is supported by the NIH (R01 AG069742 to M.B.S) and the Global Consortium for Reproductive Longevity and Equality (GCRLE-4501 to M.B.S).

## AUTHOR CONTRIBUTIONS

Conceptualization: TWPW and LEB

Methodology: TWPW, WSH, HBC, MBS, VAG, BM, DJ, KS, YS, BW, and LEB

Formal Analysis: TWPW, WSH, MR, SS, JS, CJ, SK, and JVVI

Investigation: TWPW, WSH, HBC, MR, CSB, SS, JS, HB, CJ, AC, RCK, JVVI, BM, and LEB

Original Draft: TWPW and LEB Review and Editing: all authors

Visualization: TWPW, WSH, SS, JS, HB, CJ, JVVI, BM, DJ, and LEB

Supervision and Funding Acquisition: MBS, VAG, BM, DJ, KS, YS, BW, and LEB

